# Structure-guided generative design of peptides targeting the FtsQBL divisome complex inhibit *Escherichia coli* cell division

**DOI:** 10.64898/2026.02.27.708549

**Authors:** Pauline Rémont, Xiaosong Liu, Federico Croci, Ariel Mechaly, Gouzel Karimova, Minh-Ha Nguyen, Iñaki Guijarrro, Marilyne Davi, Cécile Guillon, Constantin Bogdan Ciambur, Fabrice Agou, Alix Boucharlat, Ahmed Haouz, Jeanne Chiaravalli, Daniel Ladant, Olivier Sperandio

## Abstract

The discovery of antibiotics targeting Gram-negative bacteria remains limited in part by the difficulty of pharmacologically modulating protein–protein interactions essential for bacterial physiology. The divisome complex formed by FtsQ, FtsB and FtsL represents an attractive but challenging target, as its assembly relies on β-strand–mediated interface interactions within the bacterial periplasm. Here, we combined interpretable interface mapping using InDeep with hotspot-constrained RFdiffusion design to generate peptides targeting the FtsB-binding site of *Escherichia coli* FtsQ. The designed peptides mimic the native β-augmentation interaction and selectively engage the FtsQ interface both in bacterial cells and i*n vitro*. X-ray crystallography of one of these peptides in complex with FtsQ reveals that it accurately adopts the native binding geometry while introducing additional stabilizing interactions within a hydrophobic pocket. Complementary NMR analyses further show that optimized peptides adopt pre-organized β-hairpin conformations in solution consistent with the bound state. Several of these peptides disrupt bacterial cell division and inhibit growth in an *E. coli* strain exhibiting increased outer membrane permeability. Together, these results establish a structurally validated framework in which predictive interface analysis and generative design can be combined to target cooperative protein–protein interfaces in bacteria and provide a foundation for the development of divisome-targeting antibacterial strategies.

## Introduction

Since their discovery, antibiotics have been a major pillar in the protection of public health^1^. However, widespread misuse and a lack of chemical diversity have driven the rapid emergence of antimicrobial resistance (AMR). Without decisive action, AMR could lead to up to 10 million deaths per year worldwide ^2,3^. Some Gram-negative pathogens, such as *Escherichia coli* (Enterobacterales), are major contributors to this crisis and are classified as critical priority pathogens by the World Health Organization (WHO)^4^. Yet, no new class of antibiotics effective against Gram-negative bacteria has been approved in the past five decades, and, therefore, there is an urgent need to develop antibiotics with novel mechanisms of action that target previously unexplored cellular components.

The bacterial cell division machinery, known as the divisome, is a dynamic multiprotein complex responsible for septal peptidoglycan synthesis and constriction of the cell envelope. The divisome assembles at the bacterial septum and is composed of more than ten essential proteins that interact through a network of transient protein–protein interactions^5^. Divisome assembly is initiated by the polymerization of the cell division protein FtsZ at midcell, forming a cytoskeletal scaffold known as the Z ring. This structure is stabilized and tethered to the inner membrane by FtsA, ZipA, and Zap proteins. Subsequently, several essential membrane proteins - including FtsK, FtsQ, FtsL/FtsB, FtsW, FtsI, and FtsN - are sequentially recruited to the Z ring, leading to the activation of the core divisome complex^6^. The divisome represents a particularly attractive novel antibacterial target as it is essential for bacterial survival and it has no eukaryotic counterparts. To date, only a single antibiotic candidate targeting divisome assembly by inhibiting the initial cytosolic protein FtsZ has entered clinical trials. However, this compound is primarily effective against Gram-positive bacteria, as its access to its cytoplasmic target is severely limited in Gram-negative organisms^7^. The FtsQBL complex is a trimeric membrane assembly that plays a central role in the divisome function. Indeed, FtsQ is recruited by the chromosome segregator FtsK through its POTRA domain and then associates with FtsB and FtsL at the divisome. The trimeric FtsQBL complex then recruits of the peptidoglycan (PG) synthase complex, made of FtsW and FtsI (FtsWI), and activates it, after further recruitment of FtsN ^8^. The FtsQBL complex is highly conserved, present at low cellular abundance, and features a key protein–protein interaction interface located in the periplasm ^9–14^. The periplasmic localization of this target represents a major advantage, as it is more easily accessible to antibacterial agents (with only the outer membrane to cross)^15^. Previous studies have shown that mutations that block the assembly of the FtsQBL complex, inhibit cell division, inducing a filamentous bacterial phenotype, and ultimately leading to cell death ^9–14^. Moreover, the FtsQBL complex has already been validated as a promising target for antibacterial drug development^16^.

However, the design of protein–protein interaction (PPI) inhibitors remains particularly challenging^17–19^. Chemical libraries traditionally used for screening are often poorly suited to protein–protein interaction interfaces, which are typically larger and flatter than catalytic pockets, making them difficult to target with conventional small molecules. In this context, the recent development of deep-learning–based approaches, such as RoseTTAFold diffusion ^20^ (RFdiffusion), offers new opportunities to design peptides capable of binding to protein interfaces and thus disrupting PPIs.

In this study, we describe the design with RFdiffusion, of short peptides that are able to bind to the FtsB-binding site (FbBS) on the *Escherichia coli* FtsQ protein. They were designed as beta hairpins mimicking the two short beta strands of the two distinct partners, FtsB and FtsL, that associate to a beta strand of FtsQ. Interactions between the designed peptides and FtsQ were characterized *in vivo* by a bacterial two-hybrid (BACTH) assay, and *in vitro* by biolayer interferometry (BLI) and fluorescence polarization (FP) assays. After further *in silico* optimization, the best peptides were shown to block cell division when overproduced in bacteria. The antimicrobial activities of few synthetic peptides were then demonstrated on the *E. coli* imp4213 strain that exhibits increased outer membrane permeability. The two best peptides were shown by nuclear magnetic resonance, NMR, to adopt a beta hairpin structure in solution. The X-ray structure of one of these peptides bound to the periplasmic domain of FtsQ fully confirmed the binding mode predicted by RFdiffusion. In parallel, the peptide specificities and selectivity were established using orthologous FtsQ proteins.

Our results thus show that RFdiffusion is an efficient approach to design β-hairpin peptides that mimic protein–protein interaction interfaces. The AI-designed peptides specifically bind to the FtsB/FtsL interaction groove on FtsQ and selectively disrupt bacterial cell division and growth. They represent attractive leads to develop a new class of antimicrobials.

## Results

### Design of FtsQ Peptide Binders Using RFdiffusion

The membrane protein FtsQ acts as a central scaffold within the bacterial divisome, and associates with its binding partners FtsB and FtsL through a β-strand–mediated interface. We hypothesized that short synthetic peptides capable of reproducing this β-augmentation interaction could mimic FtsB/FtsL binding, and thus disrupt the association of FtsB/FtsL with FtsQ. To test this hypothesis, we combined the rational design of native-like peptide mimetics with AI-assisted peptide generation using RFdiffusion.

As an initial approach, three peptides were designed as direct FtsB/FtsL mimetics, corresponding to the native β-strand interface segments of FtsB and FtsL. Conservative amino acid substitutions were introduced to improve solubility and synthetic accessibility while preserving the strand register and hydrogen-bonding geometry of the native FtsQ–FtsB/FtsL complex. These peptides, designed without RFdiffusion, served as baseline mimics for subsequent comparisons.

To explore a broader structural space of FtsQ-binding peptides, we next employed RFdiffusion in binder-design mode to generate compact peptide scaffolds incorporating the key FtsB/FtsL recognition motif. In this motif-scaffolding design strategy, the β-strand element TFYRL, identified in the native FtsQ/FtsB interface by structural and InDeep-based interface analysis, was pre-positioned within the FtsQ binding groove in its native orientation thereby constraining diffusion sampling toward experimentally supported interaction geometries. RFdiffusion was then used to generate flanking backbone segments around this constrained motif, allowing the model to propose alternative peptide frameworks that preserved β-augmentation while diversifying the overall topology and sequence context.

For these initial RFdiffusion-based designs, 1,000 peptide backbones were generated, and each backbone was assigned two sequences using ProteinMPNN in complex mode, while keeping FtsQ fixed. All peptide–FtsQ complexes were evaluated using AlphaFold2. Designs were retained if they displayed a predicted interface alignment error (pAE_interaction_) below 10, showed no steric clashes, and maintained continuous β-strand pairing within the FtsQ binding groove. The remaining models were clustered based on the backbone RMSD and sequence identity to ensure structural diversity, and representative candidates were selected for experimental validation.

Eight peptides were selected for experimental validation: three rationally designed FtsB/FtsL mimetics and five RFdiffusion-derived motif-scaffolded designs (Table 1).

**Table 1.**
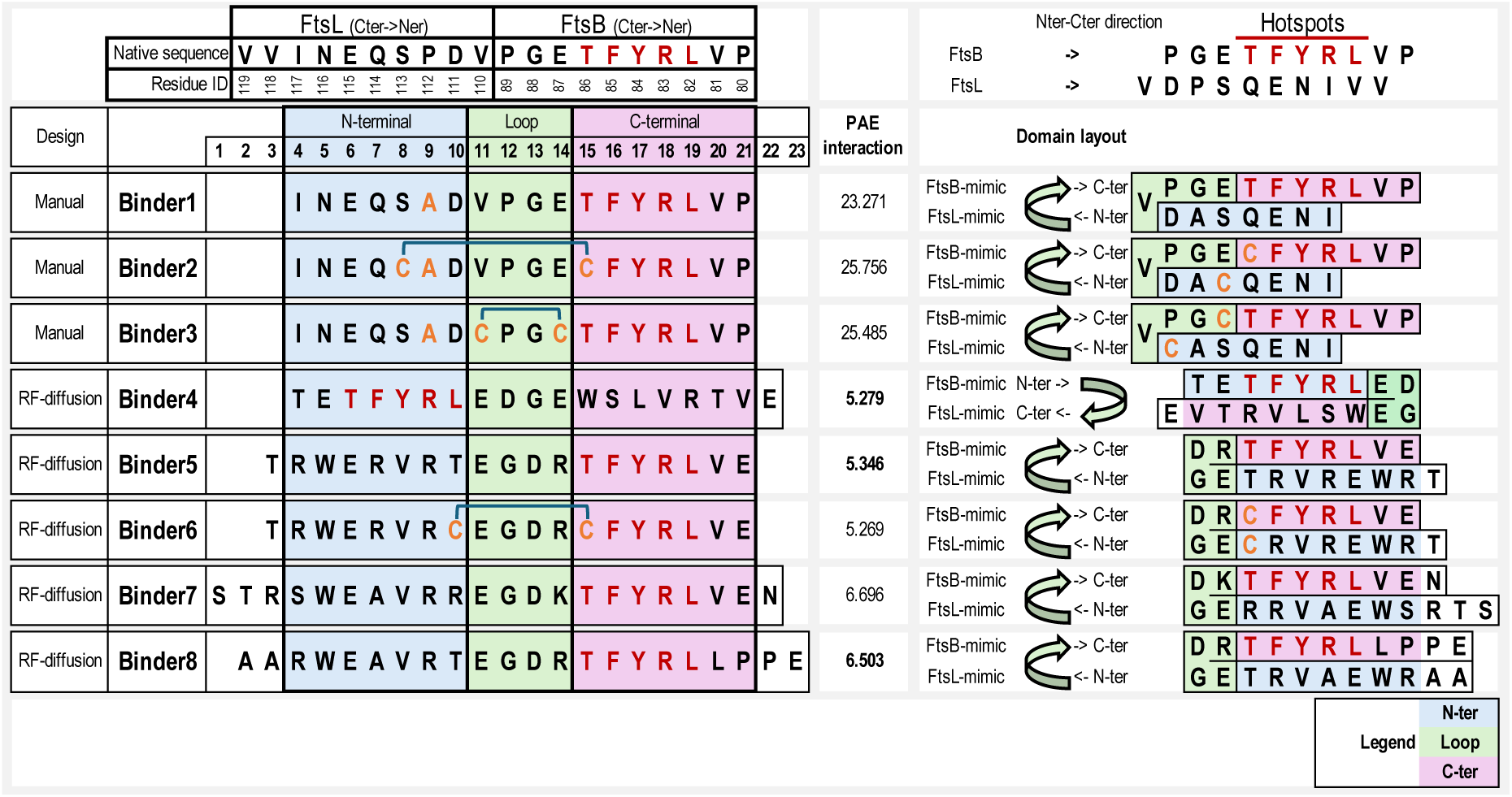
RFdiffusion-designed peptides targeting the FtsB binding site on FtsQ.

Native FtsB (PGETFYRLVP, C-terminus to N-terminus) and FtsL (VVINEQSPDV, C-terminus to N-terminus) β-strand sequences are shown at the top as reference motifs defining the FtsQ binding cassette. Below, manually designed mimetic peptides (Binders Bd1–Bd3) reproduce these native sequences with minor stabilizing substitutions, including cysteine variants. At the bottom, RFdiffusion-designed binders (Binders Bd4–Bd8) incorporate the conserved TFYRL β-strand motif within distinct scaffold contexts, featuring diverse N- and C-terminal extensions or loop insertions. The schematic on the right indicates peptide directionality (N- to C-terminus), strand orientation relative to the FtsQ binding groove, and the location of hydrophobic “hot-spot” residues that are critical for β-strand augmentation.

Sequence analysis revealed that all RFdiffusion-generated scaffolds converged toward preserving the β-strand recognition core characteristic of the native FtsQ/FtsB/FtsL interface. The native FtsB segment (PGETFYRLVP) and complementary FtsL strand (VDPSQENIVV) were used as reference templates to define the β-augmentation register within the FtsQ binding groove. The three manually designed mimetic peptides (Binders Bd1, Bd2 and Bd3) closely reproduced these native motifs, incorporating conservative substitutions or cysteine residues to enhance stability and enable potential strand pairing between FtsB and FtsL-like segments.

In contrast, the five RFdiffusion-generated designs (Binders Bd4, Bd5, Bd6, Bd7, and Bd8) diversified both peptide termini around the central TFYRL motif while preserving the hydrophobic and hydrogen-bonding patterns required for β-strand augmentation. All RFdiffusion-derived peptides retained the native strand orientation and interfacial residue composition necessary for FtsQ binding, indicating that the model preserved the essential FtsB/FtsL-derived geometry while introducing substantial sequence and topological changes (Figure 1).

**Figure 1:**
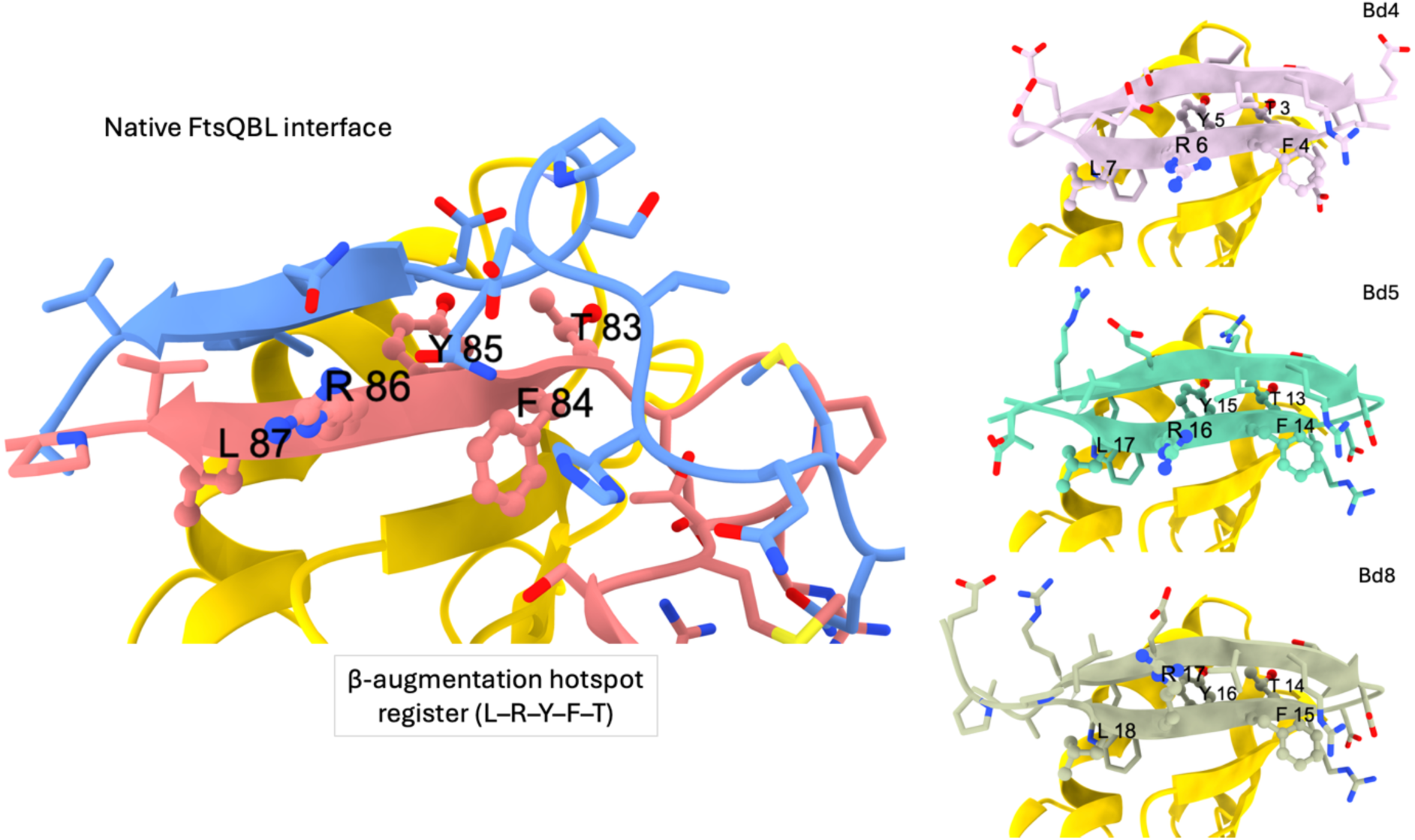
Structural models of FtsQ in complex with FtsB/FtsL or RFdiffusion-designed peptides. Structural models illustrating the binding of native and designed peptides to the periplasmic groove of E. coli FtsQ. The upper left panel shows the native FtsQ–FtsB–FtsL complex (PDB ID: 8hhh), in which the β-strands of FtsB (blue) and FtsL (pink) engage FtsQ through antiparallel β-strand augmentation, centered on the conserved T_83_F_84_Y_85_R_86_L_87_ motif. The remaining panels display the AlphaFold2-predicted complexes of FtsQ with three RFdiffusion-designed peptides: Bd4 (top right, light pink), Bd5 (middle right, turquoise), and Bd8 (bottom right, grey). All three designed peptides bind the same periplasmic groove on FtsQ, recapitulating the β-strand augmentation geometry and hydrophobic packing characteristics of the native FtsB/FtsL interface. Notably, the strand orientation differs among designs. In Bd4, the N-terminal segment adopts an FtsB-like role, whereas the C-terminal segment mimics FtsL, resulting in a topology analogous to the native FtsB–FtsL strand pairing. In contrast, in Bd5 and Bd8, the orientation is reversed: the C-terminal segment occupies the FtsB-equivalent position within the FtsQ groove, while a loop connects the N-terminal, FtsL-like region. These alternative strand orientations illustrate how RFdiffusion preserves native β-strand augmentation while sampling diverse chain directions and connectivity patterns within a conserved binding architecture.

### RFdiffusion-Designed Peptides Bind Specifically to the FtsB-Binding Site of FtsQ

To investigate the interactions between the designed peptides (Table 1) and FtsQ in bacterial cells, bacterial two-hybrid (BACTH) assays were performed either with the full-length FtsQ natively inserted in the membrane or with only the C-terminal, periplasmic domain of FtsQ expressed in the cytosol. The BACTH assay relies on the interaction-mediated reconstitution of the catalytic domain of adenylate cyclase (AC). The two complementary AC fragments, T25 and T18, were genetically fused to the N-termini of the peptides and the FtsQ variants. Hybrid proteins were co-expressed in an *E. coli cya* strain, and protein–protein interactions were quantified by measuring β-galactosidase activity.

For assays with the full-length FtsQ, the entire protein was fused to the C-terminus of T18 (pUT18C-FtsQ plasmid), while the BdX binders were fused to the T25 fragment via a synthetic transmembrane domain (OppB) in order to target the peptides to the periplasm (pKTM25-BdX plasmids). For the cytosolic assay, the BdX peptides were directly fused to the T25 fragment (pKT25-BdX plasmids), while the periplasmic part of FtsQ (residues 49–276) was fused to T18 (pUT18C-FtsQ_49–276_ plasmid).

As anticipated, the control hybrid peptides Bd1, Bd2, and Bd3, did not interact with FtsQ as indicated by the background β-galactosidase activity (i.e. similar to that of an unrelated peptide, ZIP; Figure 2A, B). In contrast, the RFdiffusion-generated binders Bd5, Bd6, Bd7, and Bd8 interacted with the full-length FtsQ inserted in the membrane (i.e. in its native state, Figure 2A) as well as with only its periplasmic domain expressed in the cytosol (Figure 2B). This latter observation indicates that the Bd5–Bd8 peptides directly interacted with the isolated periplasmic domain of FtsQ, independently of the proper insertion of FtsQ in the membrane and of other cell division components. Of note, the T25-Bd6 hybrid showed only weak complementation with T18-FtsQ_49-276_ in the cytosolic assay, which could be due to the reducing environment not favoring disulfide bond formation between the 2 Cys residues of Bd6 (Table 1).

**Figure 2:**
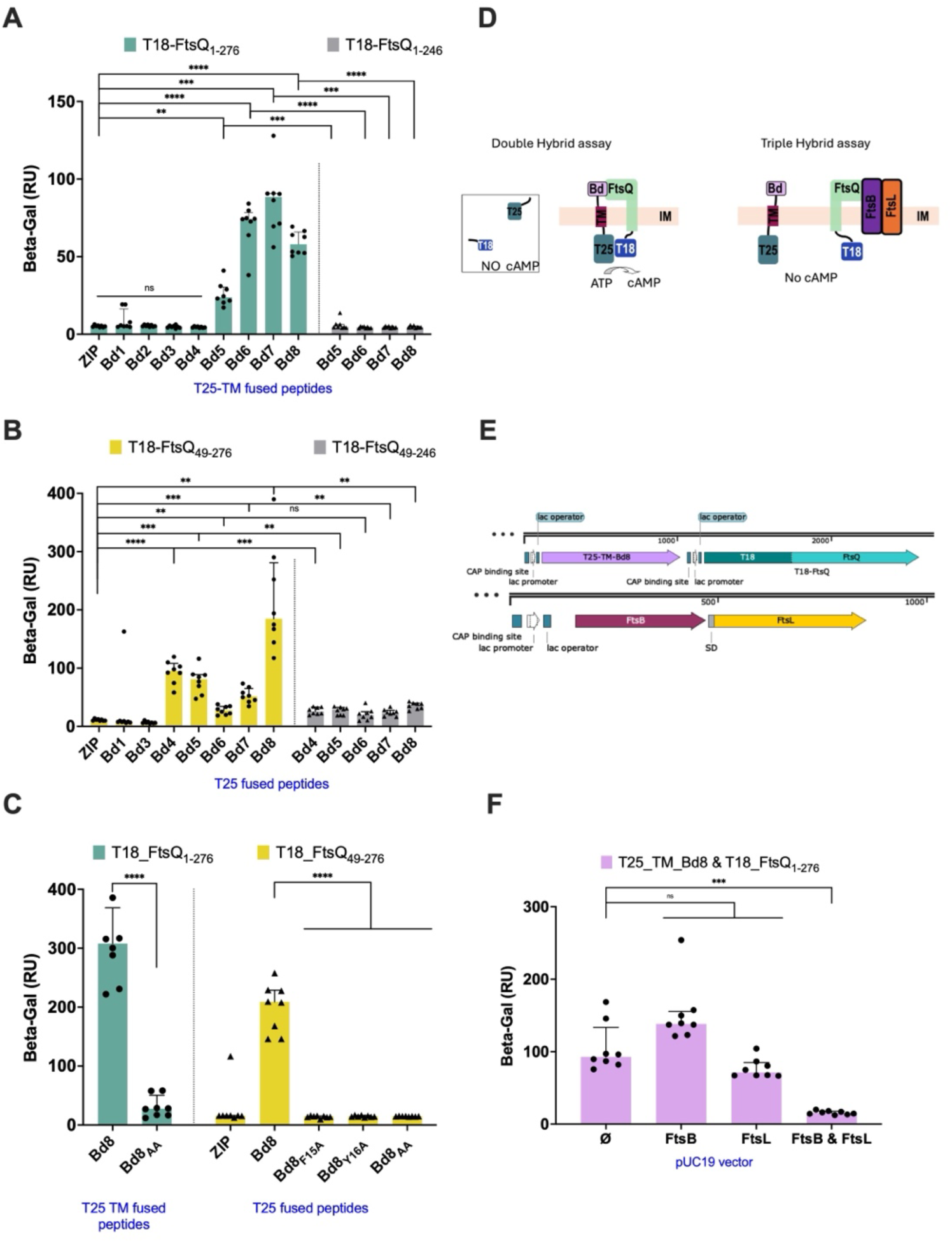
Bd peptides interaction with FtsQ proteins. Functional complementation between the indicated proteins was assessed by measuring β-galactosidase activity in liquid cultures of E. coli cya cells harboring the corresponding plasmids, as described in Materials and Methods. A leucine zipper (ZIP) motif was used as a negative control. Each bar represents the median of at least six independent cultures (technical replicates), and the error bars indicate the interquartile range. Brown-Forsythe and Welch One-way Anova test and Dunett T3 post hoc, ns > 0.05, **P < 0.01, and ***P < 0.001. **(A)** BACTH assay showing complementation between T25–transmembrane (TM) peptide fusions (pKTM25) and either full-length T18–FtsQ (residues 1–276; pUT18C-FtsQ1–276, blue) or a C-terminally truncated variant (residues 1–246; pUT18C-FtsQ1–246, grey) in E. coli DHM1 cells. **(B)** BACTH assay between T25–peptide fusions (pKT25) and either the periplasmic domain of FtsQ (residues 49–276; pUT18C-FtsQ49–276, yellow) or a C-terminally truncated version (residues 49–246; pUT18C-FtsQ49–246, gray) in E. coli DHT1 cells. The Bd2 peptide could not be cloned into the pKT25 vector and was therefore not assessed in this cytosolic assay **(C)** BACTH assays in the cytosol and periplasm assessing interactions between Bd8 variants (Bd8_F15A_, Bd8_Y16A_, Bd8_F15A,Y16A_ =Bd8_AA_) and either full-length FtsQ (pUT18C-FtsQ_1–276_, blue; in E. coli DHM1) or the periplasmic domain of FtsQ (pUT18C-FtsQ_49–276_, yellow; in E. coli DHT1). **(D)** Schematic representation of the BACTH assay in the absence (left) or in the presence (right) of co-expressed FtsB and FtsL **(E)** Schematic maps of the pKTM25-Bd8_T18-FtsQ_1–276_ and pUC19-FtsB/FtsL plasmids, showing the locations of the encoded genes, the Shine–Dalgarno sequence (SD), and the lac promote**(F)** Three-hybrid competition experiments examining interactions between T18–FtsQ_1–276_ and T25–TM–Bd8, alone (Ø) or in the presence of co-expressed FtsB, FtsL, or both FtsB and FtsL in E. coli DHM1 cells.

Interestingly, the RFdiffusion-designed peptide Bd4 interacted with T18-FtsQ_49–276_ in the cytosol, but no complementation was detected with the full-length T18-FtsQ. One possible explanation is that the fusion of the Bd4 N-terminus to the linker and synthetic transmembrane segment somehow interferes with its interaction with full-length FtsQ. Consistent with this interpretation, Bd4 adopts a predicted orientation opposite to that of the other Bd peptides, positioning its transmembrane fusion on the opposite side of the FtsQ binding groove (Figure 1). Alternatively, the native FtsB and FtsL proteins may compete more efficiently with Bd4 binding to full-length FtsQ in the periplasm.

Previous studies have established that the key interaction between FtsB and FtsQ involves their respective C termini. In addition, Kureisaite-Ciziene et al. showed that single point mutations in either FtsQ (e.g., Y248K) or FtsB (e.g., F84A or Y85A) induce bacterial filamentation ^11^, highlighting the functional importance of this interface. To confirm that the designed Bd peptides specifically bind to the C-terminal region of FtsQ, we performed three complementary experiments.

First, we used truncated FtsQ variants lacking the C-terminal region (FtsQ_1–246_ or FtsQ_49–246_) in BACTH assays. Deletion of the FtsQ C-terminus abolished complementation between the hybrid peptides and FtsQ, indicating that the C-terminal extremity of FtsQ that harbors the FtsB-binding β-strand is critical for interaction with the Bdpeptides (Figure 2A, B).

Second, the key aromatic residues F84 and Y85, of FtsB are conserved in the central motif (TFYRL) in all Bd peptides, we introduced alanine substitutions at the corresponding positions in the Bd8 peptide sequence: Y16A (Bd8_Y16A_), F15A (Bd8_F15A_), and double mutant F15A/Y16A (Bd8_F15A/Y16A_ = Bd8_AA_). Substitution of either aromatic residue with an alanine resulted in a complete loss of interaction between the Bd8 peptide variants and the T18–FtsQ_49–276_ construct (Figure 2C). These results indicate that the aromatic residues F15 and Y16 of Bd8 are critical for binding to FtsQ, consistent with their functional equivalence to the F84 and Y85 anchor residues of FtsB, respectively.

Together, these results suggest that the designed peptides specifically target the FtsB-binding site on FtsQ. To further confirm this, we examined the effect of co-expressing FtsB and/or FtsL on the complementation between T18–FtsQ and T25–TM–Bd8 hybrids using a three-hybrid BACTH assay (Figure 2D). The T18–FtsQ coding sequence was cloned into pKTM25-Bd8 to generate pKTM25-Bd8_T18-FtsQ, a plasmid that co-expresses both T18–FtsQ and T25–TM–Bd8 hybrids. This plasmid was then co-transformed in DHM1 with pUC19 derivatives, expressing either FtsB (pUC19-FtsB), FtsL (pUC19-FtsL), or both proteins simultaneously (pUC19-FtsB-FtsL) (Figure 2E). Overexpression of both FtsB and FtsL resulted in a complete loss of interaction between the T25–TM–Bd8 and T18–FtsQ hybrid (Figure 2F). In contrast, overproduction of FtsB or FtsL alone did not significantly affect Bd8’s binding to FtsQ as compared to the empty pUC19. These results suggest that the FtsB/FtsL dimeric complex, but not either protein alone, effectively competes with Bd8 for binding to FtsQ. Overall, these data confirm that Bd8 and the native FtsB/FtsL complex compete for the same binding site on FtsQ.

### Biophysical Characterization of Synthetic Peptide Binding to the Periplasmic Domain of FtsQ

Biolayer interferometry (BLI) and fluorescence polarization (FP) experiments were then performed to characterize the Bd peptide/FtsQ interactions *in vitro*. These experiments were carried out with the His-tagged purified periplasmic domain of FtsQ (6His-FtsQ_49–276_, overexpressed in *E .coli*, see “Material and Methods”) and with synthetic peptides, Bd4, Bd5, and Bd8, which were selected based on their strong interaction signals observed in the cytosolic BACTH assays (Figure 2B). Synthetic peptides Bd1 and Bd8_F15A_, carrying the alanine-substituted Bd8 variant, were also characterized as negative controls.

Biolayer interferometry (BLI) assays were performed using biotinylated peptides (Bd1, Bd4, Bd5, and Bd8) synthesized by Genosphere and immobilized on streptavidin biosensors (Sartorius, Germany). The purified periplasmic domain of FtsQ (6His-FtsQ_49–276_) was used as an analyte (Figure S1). Peptides Bd4, Bd5, and Bd8 exhibited strong binding signals to FtsQ, whereas Bd1 did not show any detectable interactions. Although the peptides are supposed to interact with FtsQ according to a 1:1 binding model, the experimental data were better described by a 2:1 heterogeneous ligand binding model (Figure 3A). This behavior is consistent with previous reports describing the interaction between FtsQ and a FtsB-derived peptides^21^. Fitting the data using the 2:1 model indicated that peptides Bd5 and Bd8 bind to FtsQ with two apparent affinity components, with dissociation constants of approximately 0.3–0.4 µM (K_D1_) and 1.3–1.4 µM (K_D2_). Peptide Bd4 also bound FtsQ, with a K_D1_ of approximately 0.3 µM and a lower-affinity component with a K_D2_ of approximately 4.1 µM. (Figure 3B).

**Figure 3:**
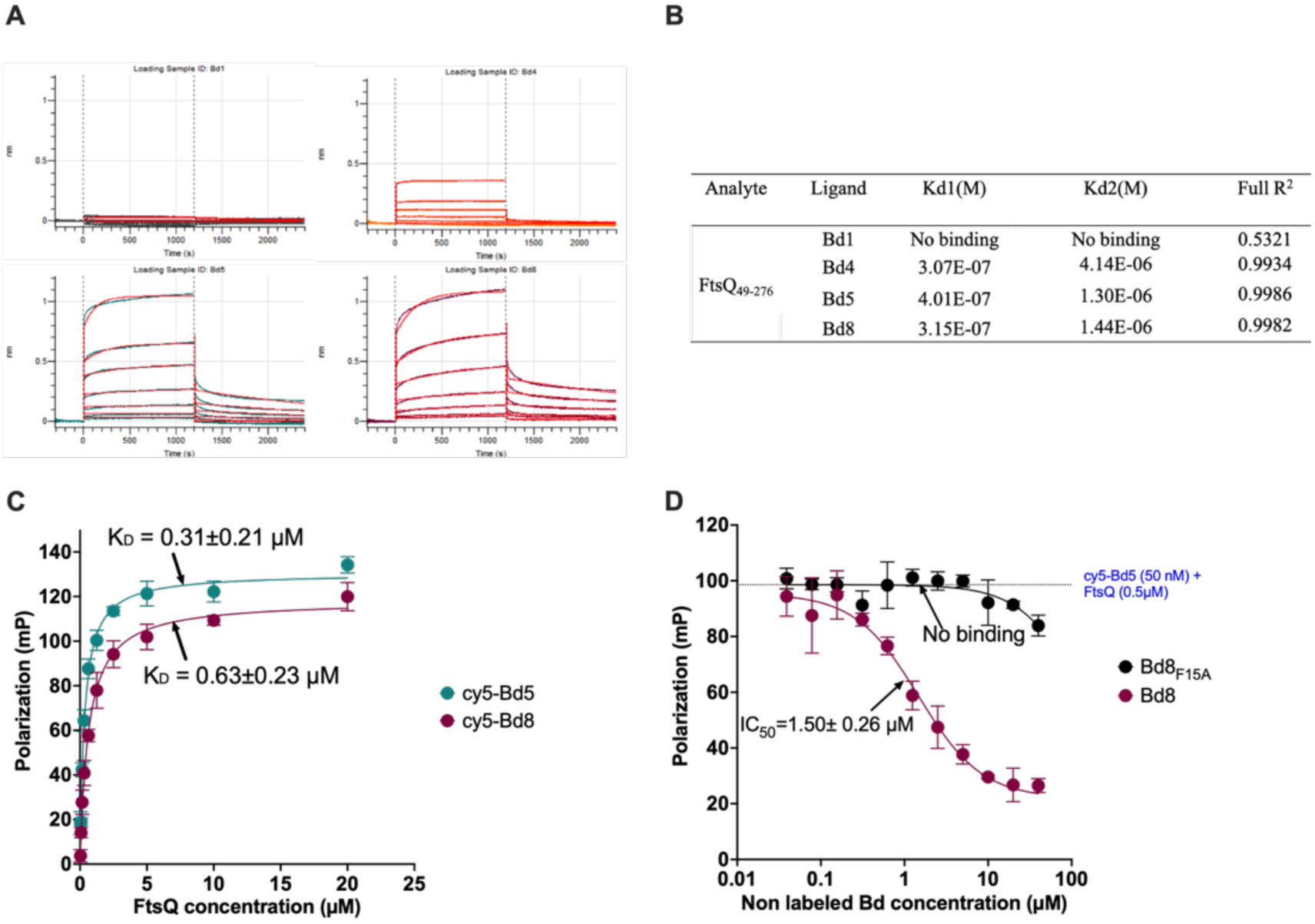
In vitro analysis of Bd peptide binding to FtsQ **(A)** Biolayer interferometry (BLI) association–dissociation curves for interactions between the periplasmic domain of FtsQ and the indicated biotinylated peptides immobilized on streptavidin-coated biosensors (Bd1, top left; Bd4, top right; Bd5, bottom left; Bd8, bottom right). For each peptide, eight different concentrations of analyte (6His–FtsQ_49–276_) were tested. Data were analyzed using Octet Analysis Software and fitted with a 2:1 heterogeneous ligand binding model; red curves indicate the fitted traces. **(B)** Dissociation constants (K_D_ in M) for each peptide calculated from BLI data. **(C)** Fluorescence polarization (FP) assay of binding of fluorescently labeled peptides Cy5–Bd5 (blue) or Cy5–Bd8 (purple) (both at 50 nM) to the 6His tagged periplasmic domain of FtsQ (0–20 µM). Data points represent the means of three independent measurements, and error bars indicate the standard deviation. The data were fitted using a one-site saturation binding model in Prism software. **(D)** FP competition assays measuring the displacement of Cy5–Bd5 by increasing concentrations of unlabeled Bd8 (purple) or Bd8_F15A_ (black). Assays were performed using 50 nM Cy5–Bd5 and 0.5 µM 6His–FtsQ_49–276_. Data points represent the mean of three independent measurements with standard deviation and were fitted using a variable-slope dose–response inhibition model in Prism software. For all FP experiments, the curves depict one representative assay among three independent assays, and the calculated IC_50_/ K_D_ are the means of the three independent experiments

In parallel, these interactions were validated in solution using fluorescence polarization (FP) assays with fluorescently labeled peptides (Cy5–Bd8 and Cy5–Bd5). FP measurements were consistent with the BLI results, yielding dissociation constants of 0.63 µM for Cy5–Bd8 and 0.31 µM for Cy5–Bd5, respectively (Figure 3C). Similar to the BLI assay, FP competition experiments confirmed that non-fluorescent Biotin–Bd4, Biotin–Bd5, and Biotin–Bd8 effectively displaced the fluorescent peptides, whereas the Biotin–Bd1 peptide did not compete for binding (Figure S2). Finally, FP competition assays using untagged peptides Bd8 and Bd8_F15A_ confirmed that the alanine-substituted Bd8 mutant (Bd8_F15A_) was unable to interact with FtsQ whereas Bd8 had an IC_50_ of 1.50 µM. (Figure 3D).

### Peptide optimization

To further improve the binding affinity, we performed a second round of optimization using RFdiffusion, starting from the Bd4, Bd5, and Bd8 sequences. Ten optimized sequences were selected for each initial binder (Table S1). These optimized peptides were first screened using the BACTH approach in *E. coli DHT1* cells. The best ones were tested in a second assay performed in the *E. coli* DHM1 strain under more stringent conditions (i.e.in the absence of IPTG) (Table S2). This was done to reduce the expression levels of the hybrid proteins and favor the detection of higher-affinity interactions. Under these conditions, several T25-hybrid peptides exhibited significantly stronger interaction signals with T18–FtsQ_49–276_ than their corresponding parental peptides (Figure 4A, Table S2).

**Figure 4.**
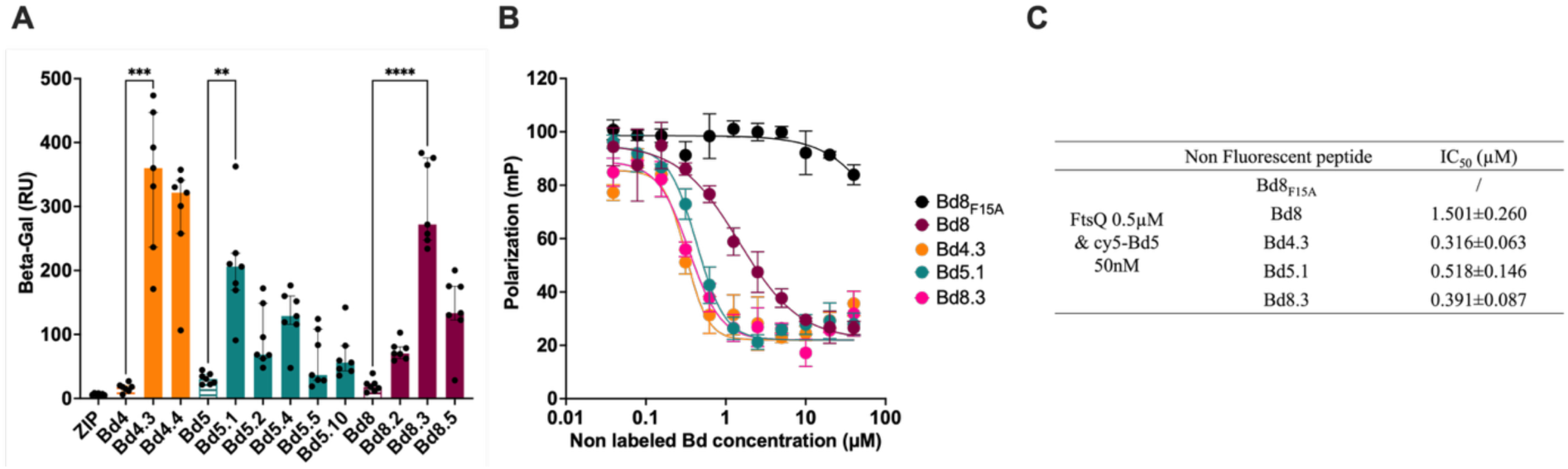
Characterization of the optimized peptides. **(A)** Bacterial two-hybrid (BACTH) assays assessing functional complementation between the periplasmic domain of FtsQ (pUT18C-FtsQ_49–276_) and a series of optimized Bd peptides fused to T25 (pKT25-BdX). Optimized derivatives of Bd4 (orange), Bd5 (blue), and Bd8 (purple) were tested in E. coli DHM1 cells under stringent conditions (i.e. in the absence of IPTG), as described in the Materials and Methods. β-Galactosidase activity was measured in suspensions of chloroform-treated cells. Each bar represents the median of at least five independent cultures, and the error bars indicate the interquartile range. The leucine zipper peptide (ZIP) was used as a negative control. **(B)** Fluorescence polarization (FP) competition assays measuring the displacement of Cy5–Bd5 by increasing concentrations of unlabeled optimized peptides (Bd4.3, orange; Bd5.1, blue; Bd8.3, pink), the parental Bd8 peptide (purple), or the Bd8_F15A_ variant (black). Assays were performed using 50 nM Cy5–Bd5 and 0.5 µM 6His–FtsQ_49–276_. Data points represent the mean of three independent measurements, with error bars indicating the standard deviation. The data were fitted using a variable-slope dose–response inhibition model in Prism software. The curves depict one representative assay among three independent ones. **(C)** Half-maximal inhibitory concentrations (IC_50_) for each peptide derived from the mean of three independent FP competition experiments.

These results were further validated *in vitro* using FP analysis. Synthetic peptides corresponding to the optimized sequences Bd4.3, Bd5.1, and Bd8.3 displayed reduced IC_50_ values, with a 3-to-5-fold decrease compared with that of the original Bd8 peptide (Figure 4B, C). In parallel, BACTH assays confirmed that the optimized T25–TM–Bd5.1 and T25–TM–Bd8.3 peptides specifically interacted with T18–FtsQ (i.e. in native membrane environement) but not with T18–FtsQ_1–246_ lacking the critical C-terminal FtsB-binding β-strand (Figure S3). In addition, as observed with the parental Bd4 peptide, the optimized T25–TM–Bd4.3 peptide also did not produce a detectable interaction signal with T18–FtsQ (Figure S3).

### Effect of FtsQ-binding peptides on bacterial phenotype and growth

To further assess the impact of FtsQ-binding peptides on *E. coli* (MG1655) cell division, Bd5, Bd8, Bd4.3, Bd5.1, and Bd8.3 were overproduced as fusions to the T18 fragment, followed by a transmembrane (TM) domain to target the peptides to the periplasm (see Materials and Methods). BACTH assays confirmed that these hybrid peptides, except Bd4.3, retained their ability to interact with FtsQ (Figure S3).

Microscopic analysis of bacterial morphology following Bd peptide overproduction revealed distinct phenotypic effects. Whereas overproduction of the original peptides Bd5 and Bd8 did not noticeably affect cell morphology, overproduction of the optimized peptides Bd5.1 and Bd8.3 significantly interfered with bacterial cell division. In particular, Bd5.1 overproduction induced pronounced bacterial filamentation and was associated with a marked impairment of bacterial growth (Figure 5). This filamentous phenotype is consistent with the disruption of divisome assembly, likely through interference of FtsQ association with FtsB/FtsL. Importantly, when FtsQ was overproduced (from a compatible, low copy plasmid) in the same cells, overexpression of the Bd5.1 peptide did not any more trigger detectable bacterial filamentation (Figure 5A, B). The rescue by FtsQ of the Bd5.1-induced disruption of divisome assembly strongly suggest that Bd5.1 acted as an FtsQ inhibitors by blocking further association with FtsB/FtsL. It should be noted also that peptide Bd4.3 did not induce detectable morphological changes, in agreement with previous BACTH experiments showing that the membrane-tethered Bd4.3 fusion cannot interact with full-length FtsQ (Figure S3, Figure S4).

**Figure 5:**
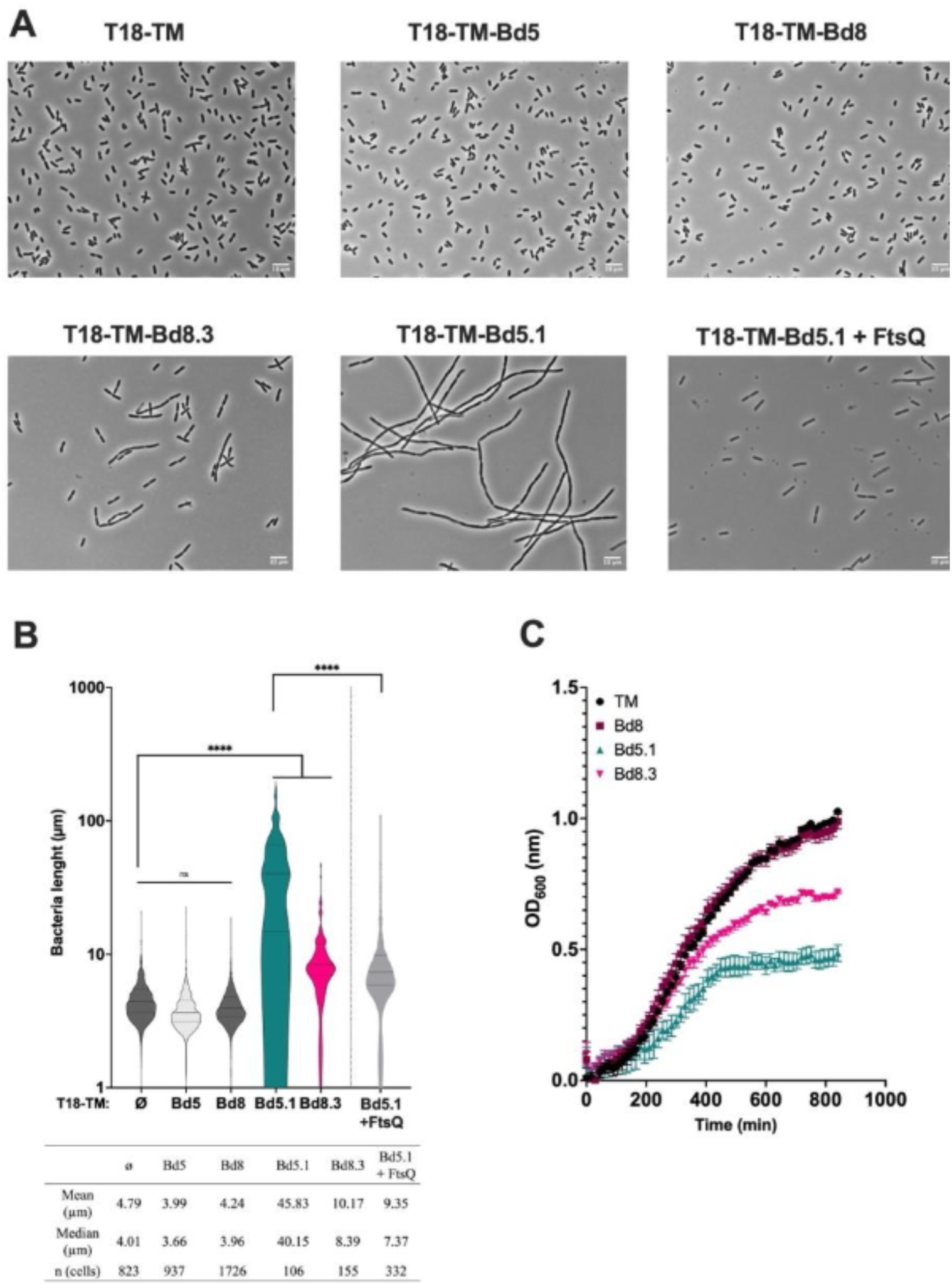
Impact of peptide overproduction on bacterial division **(A)** Phase-contrast microscopy images of E. coli MG1655 cells overproducing the indicated T18–TM–Bd peptides (plasmid pUTM18C-BdX) after 6 h 30 min of induction with 0.5 mM IPTG). Images were acquired at ×100 magnification. Scale bar 10µm **(B)** Quantification of bacterial cell length following 6 h 30 min of peptide overexpression, measured using ObjectJ software. Median and interquartile range are showed. One way Brown-Forsythe and Welch Anova test with Games-Howell Post hoc, ns > 0.05, and ****P < 0.0001 **(C)** Growth monitoring of E. coli MG1655 cultures overexpressing T18–TM–Bd peptides, assessed by optical density at 600 nm (OD_600_). Dots represent the mean of three independent cultures (technical replicates) and error bars the standard deviation.

In a second set of experiments, we directly assessed the ability of synthetic peptides to modulate bacterial growth. To this end, we used the *E. coli* lptD4213 (imp) strain, which exhibits increased outer-membrane permeability to large compounds due to a mutation in the *lptD* gene^22^. We first tested the optimized peptides Bd4.3, Bd5.1, and Bd8.3. While Bd4.3 displayed a minimum inhibitory concentration (MIC) of 25 µM and led to a filamentous bacterial phenotype, neither Bd5.1 nor Bd8.3 affected the growth of *E. coli* lptD4213 at concentrations up to 50 µM (Table 2, Figure 6). Peptide Bd5.1 was able to induce few filamentous cells at 100µM after 2h30 of exposure but without significant impact on growth.

**Figure 6:**
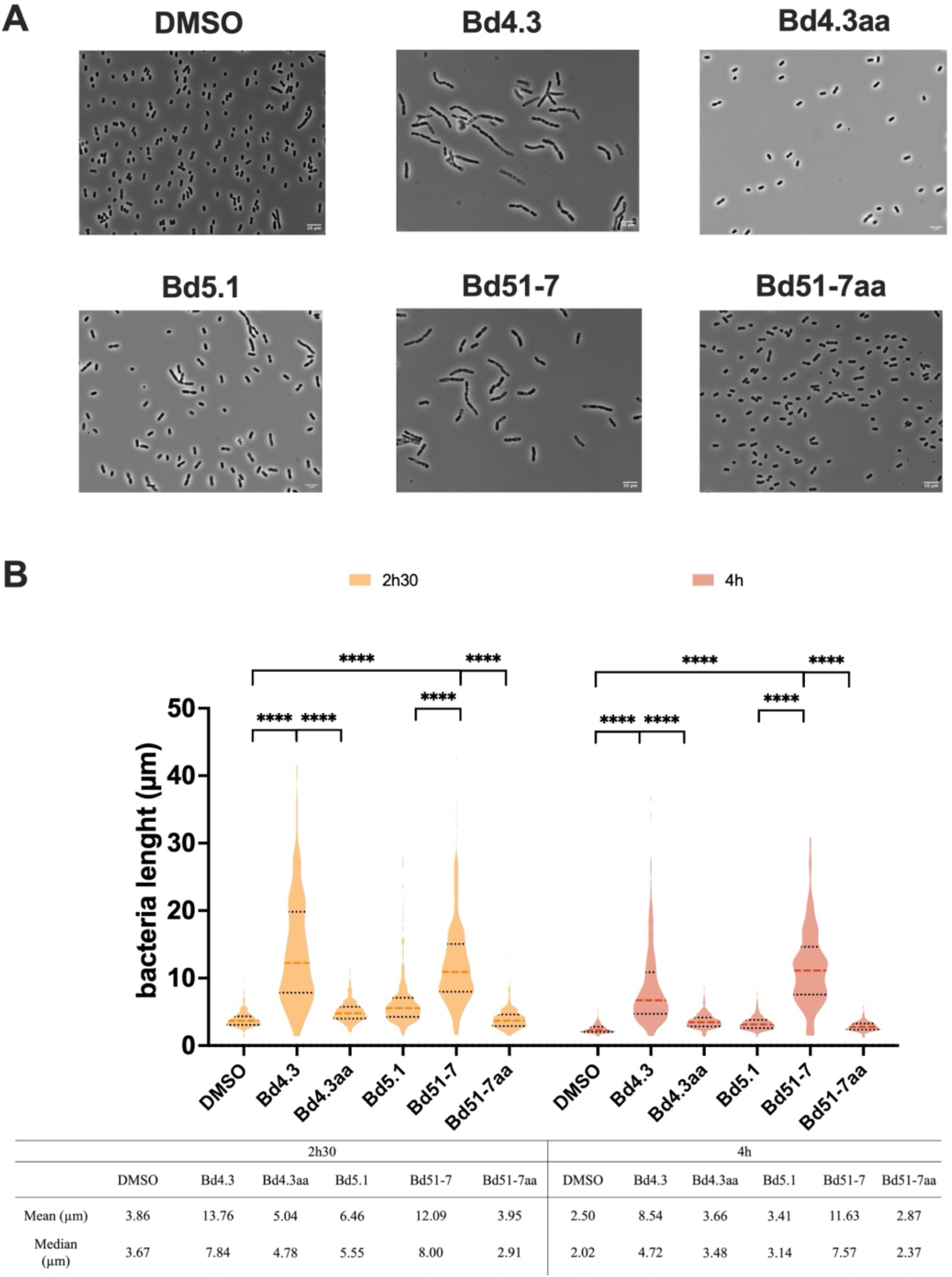
E. coli lptD4213 morphological change induced by Bd peptides **(A)** Phase-contrast microscopy images of E. coli lptD4213 in the presence of either DMSO, peptide Bd5.1 (100µM), peptides Bd4.3 and Bd4.3aa (50µM) or peptides Bd51-7 and Bd51-7aa (12.5µM) after 2h30. Peptides Bd4.3 and Bd51-7 were tested at a concentration equal to 2 x MIC as well as their corresponding alanine variants. Images were acquired at ×100 magnification. Scale bar 10µm **(B)** The violin plot shows the corresponding cell length distribution of samples treated with either DMSO or Bd4.3 (50µM) or Bd51-7 (12.5µM) or Bd51-7aa (12.5µM) after 2h30 and 4 h. Median and interquartile range are represented. One way Brown-Forsythe and Welch Anova test with Games-Howell Post hoc, ****P < 0.0001

**Table 2:**
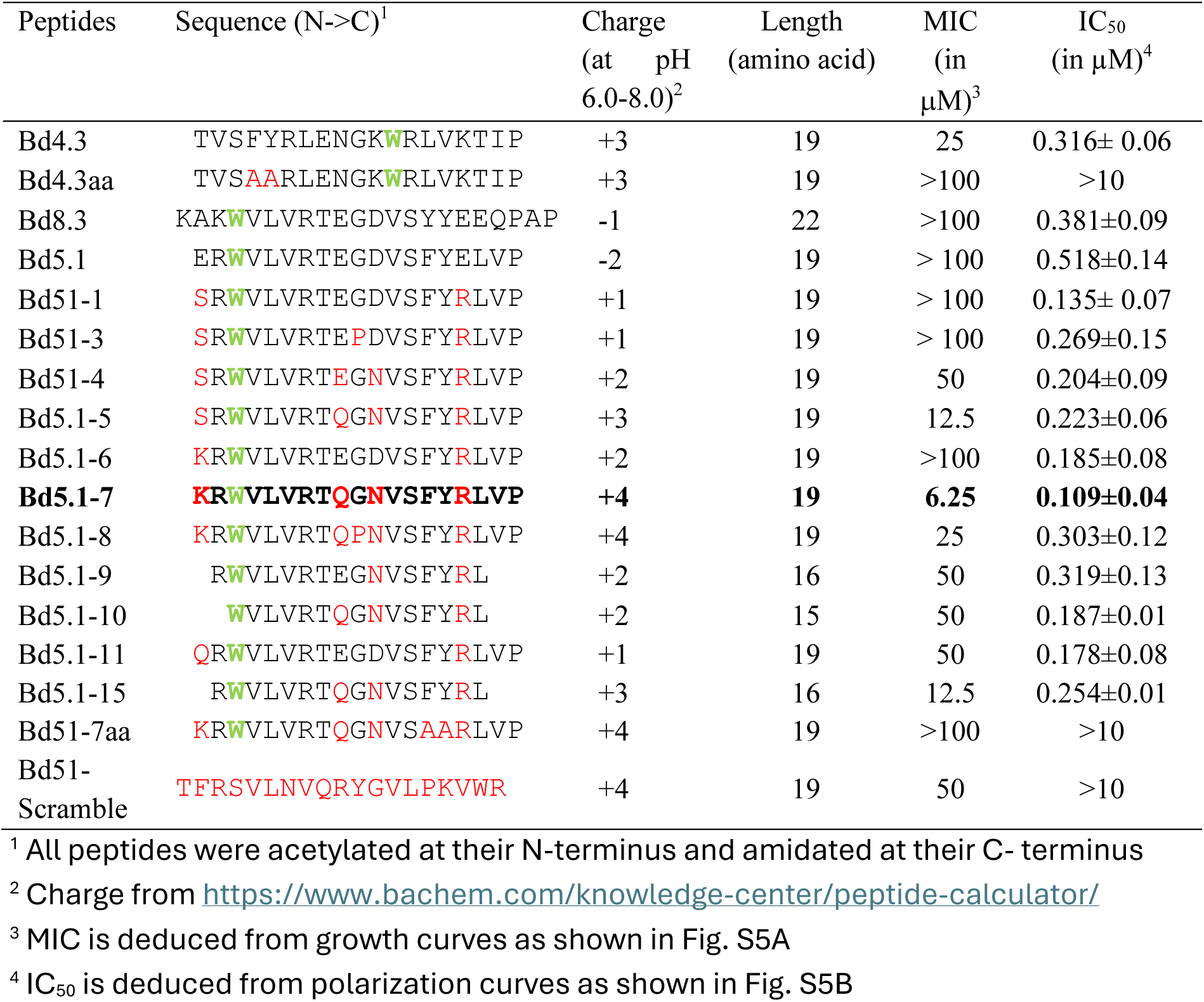
Effect of synthetic FtsQ-binding peptides on *E. coli lptD*4213 bacterial growth

Because peptides Bd5.1 and Bd8.3 are negatively charged at neutral pH (net charge = −2), we hypothesized that they were unable to effectively access the periplasm of *E. coli*. We therefore designed and synthesized additional Bd5.1 derivatives in which several residues were modified to reverse the overall charge. Several of these positively charged derivatives inhibited the growth of *E. coli* lptD4213 (Table 2). Among them, peptide Bd5.1-7 was the most potent, inducing complete growth inhibition at 6.25 µM and a significant growth delay at 3.1 µM (Table 2, Figure S5).

In addition, after 2 h 30 min of treatment, bacteria exposed to Bd5.1-7 exhibited an elongated phenotype with multiple constrictions per cell similarly to Bd4.3, whereas DMSO-treated bacteria displayed a normal, short-cell morphology. Such chaining and elongation phenotypes have been previously associated with inhibition of bacterial cell division (Figure 6). Furthermore, fluorescence polarization assays revealed that Bd5.1-7 exhibited increased affinity for FtsQ, with an IC_50_ of approximately 0.1 µM, suggesting that charge optimization not only facilitates bacterial entry but also enhances Bd peptide/FtsQ interactions (Table 2).

Finally, the most active peptides were evaluated for cytotoxicity against MRC5 human lung fibroblast cells at a concentration of 50 µM. None of the tested peptides displayed detectable cytotoxicity, with cell viability remaining above 95% after 48 h of culture (Figure S6).

In parallel, synthetic peptides Bd4.3_AA_ (F4A and Y5A), Bd5.1-7_AA_ (F14A and Y15A), carrying aromatic to alanine substitutions, as well as a scrambled control peptide Bd5.1_-scramble_, (composed of the same amino acids as Bd5.1-7 but in a randomized sequence), exhibited no, or strongly reduced, effects on bacterial growth (Table 2, Figure S5). None of these control peptides showed detectable interaction with FtsQ in fluorescence polarization assays (Figure S5B). These results indicate that the antibacterial activities of Bd4.3 and Bd5.1-7 are primarily dependent on their specific interaction with FtsQ. To directly compare the cellular effects of Bd5.1-7 with those of Bd5.1, the T18–TM–Bd5.1-7 hybrid peptide was overproduced in *E. coli* MG1655. Similar to T18–TM–Bd5.1, overproduction of T18–TM–Bd5.1-7 induced a pronounced elongated phenotype further supporting that Bd5.1-7 toxicity results from specific targeting of FtsQ (Figure S7).

### Structural characterization of Bd4.3 and Bd5.1-7 by Xray crystallography and NMR

To structurally validate the binding mode of the designed peptides, we determined the crystal structure of Bd4.3 in complex with FtsQ (Figure 7). The structure reveals that Bd4.3 binds at the same interface region engaged by the native FtsB/FtsL partners in the FtsQBL complex (PDB: 8HHH), reproducing the β-strand augmentation characteristic of the divisome assembly interface. The peptide aligns along the last FtsQ β-sheet and engages the hotspot region defined by FtsB residues Thr83, Phe84, Tyr85, Arg86 and Leu87, which were explicitly used as positional constraints during RFdiffusion-guided design. Clear and continuous electron density unambiguously supports the modeled conformation of the Bd4.3 peptide at the interface. While the overall binding geometry closely mimics the native interaction, the designed peptides systematically introduce a tryptophan residue (at position 12 for Bd4.3 or at position 3 for Bd5.1-7) that is absent from the corresponding regions of FtsB and FtsL. In the Bd4.3 structure, this residue inserts into a hydrophobic pocket adjacent to the native interface, generating additional packing interactions not exploited in the natural complex (Figure S8). These observations indicate that the design strategy successfully combined interface mimicry with targeted pocket engagement, providing a structural basis for the improved binding properties observed for this peptide series.

**Figure 7.**
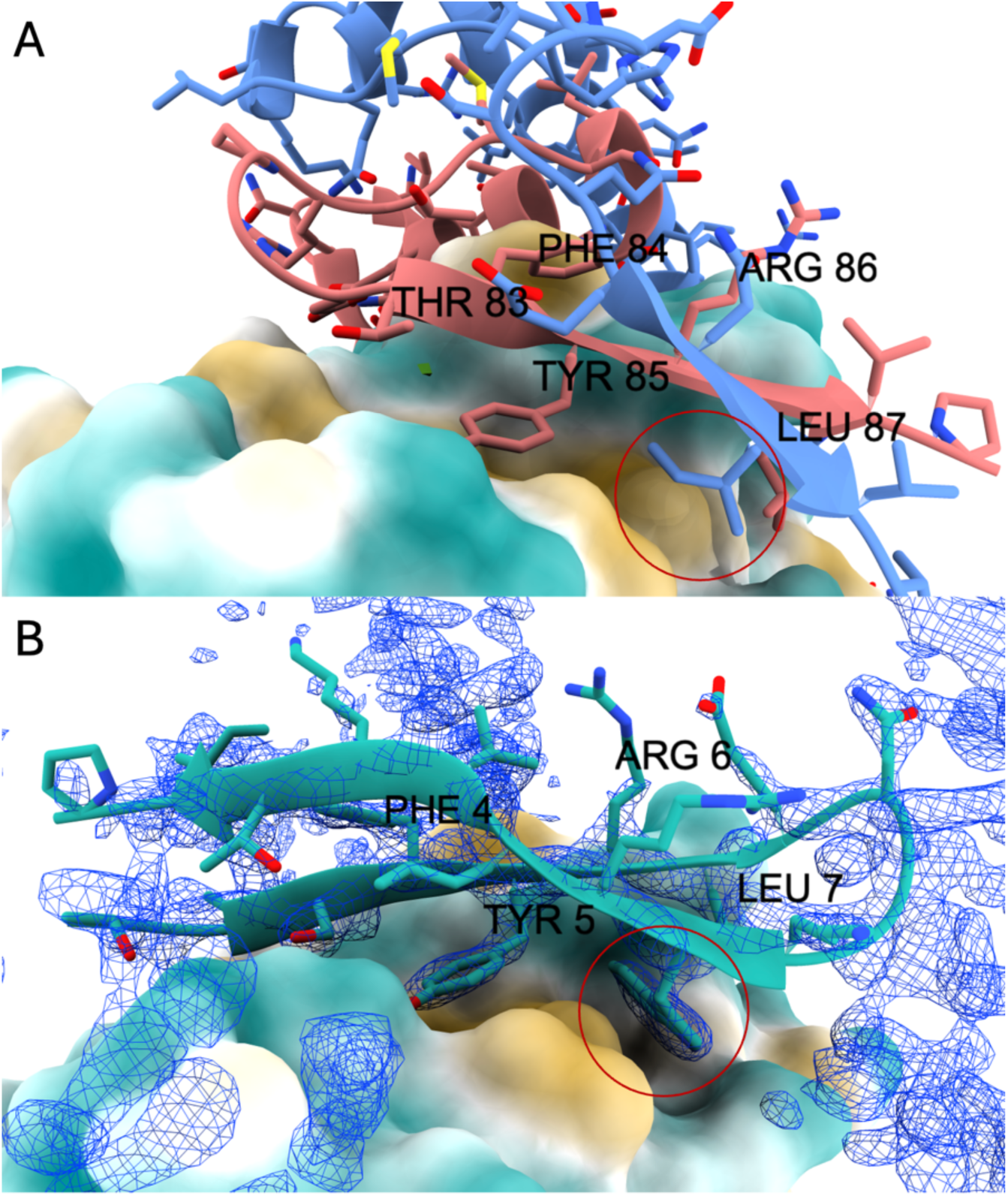
Structural comparison between the native FtsQBL interface and the designed peptide Bd4.3 bound to FtsQ. (A) Structure of the native FtsQBL complex (PDB: 8HHH) showing the β-strand interface formed by FtsB/FtsL on FtsQ. Interface hotspot residues on FtsQ (Thr83, Phe84, Tyr85, Arg86 and Leu87), used as positional constraints during RFdiffusion-guided peptide design, are highlighted. The region corresponding to the hydrophobic pocket targeted during design is indicated. (B) Crystal structure of the designed peptide Bd4.3 bound to FtsQ. Experimental electron density (blue mesh) confirms the positioning of the peptide at the targeted interface and the preservation of the β-strand augmentation geometry. A conserved tryptophan residue (position 12 in Bd4.3) present in all designed sequences (red circle), but absent from the native FtsB and FtsL sequences, inserts into a hydrophobic pocket defined by the hotspot region, providing an additional stabilizing interaction exploited during design.

The solution structures of peptides Bd4.3 and Bd5.1-7 were also investigated by NMR. ^1^H, ^13^C and ^15^N assignments were obtained in natural abundance and are summarized in Supplementary tables (TableS3 and TableS4). Secondary structure propensities and order parameters reflecting backbone dynamics on the ps-ns time range were calculated from the backbone Hα, N, CA and CB chemical shifts (Figure 8). For both peptides, the secondary structure propensities correspond to an antiparallel β-hairpin with two β-strands separated by a loop. In addition, both peptides show restricted motions on the ns-ps time range as evidenced by high order parameters (S^2^ < 0.65) of most of the backbone ^15^N nuclei, further indicating that the peptides are structured. For Bd4.3, however, signal broadening reveals exchange between different conformations on the µs-ms time range. Broadening and hence µs-ms dynamics, are indeed observed for residues Y5, R6, L7, I18 and P19.

**Figure 8.**
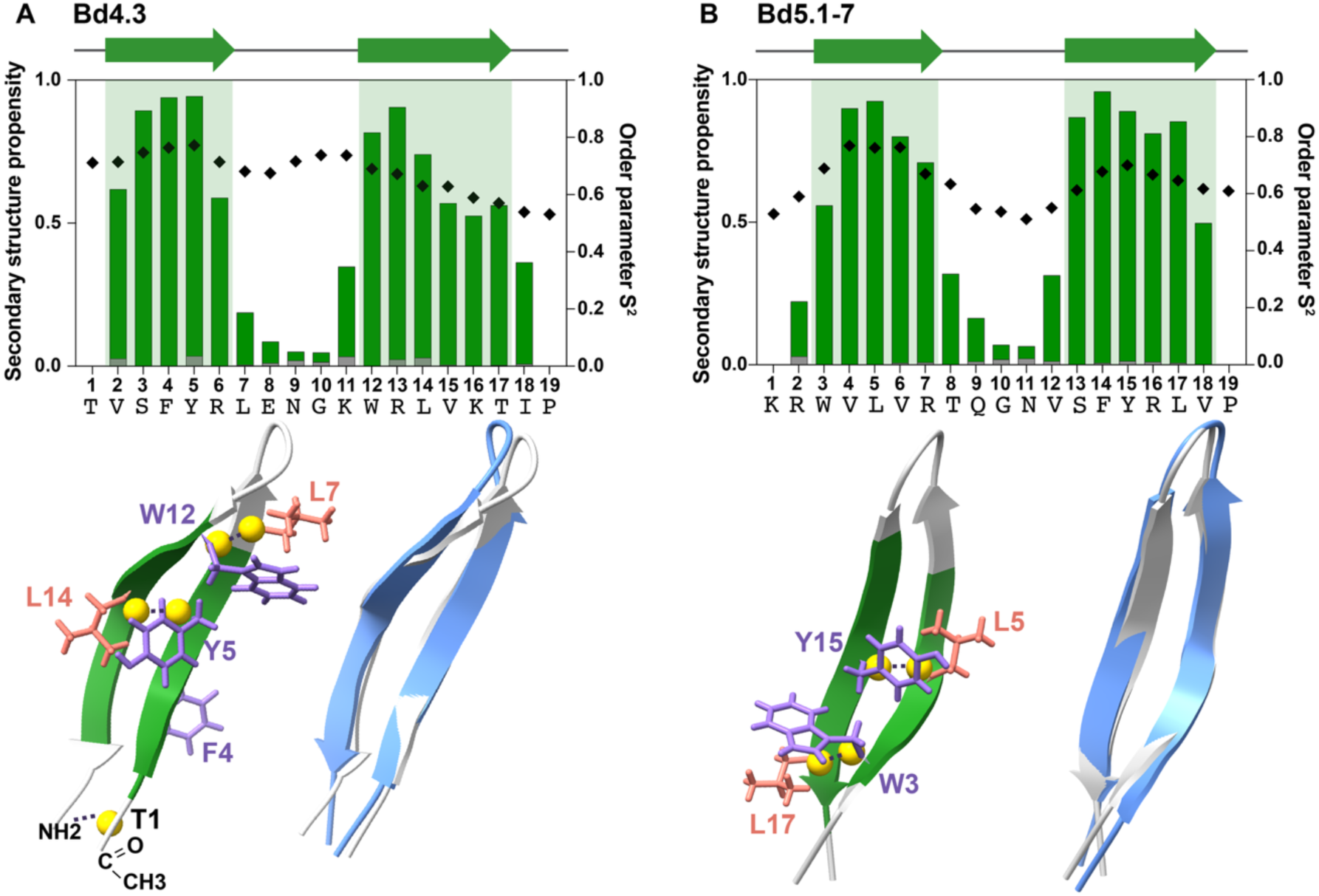
NMR solution structure of peptides Bd4.3 (A) and Bd5.1-7 (B). **Top:** ^15^N S order parameters (black diamonds) reflecting backbone dynamics on the ps-ns time and bar-chart graph of β-sheet (green) and ***α***-helix (grey) propensities plotted as a function of the residue. Regions with β-sheet conformation are highlighted in light green and the predicted strands are displayed as green arrows on the top. **Bottom**: cartoon representation of the solution structures of the peptides (left of each panel) highlighting the aromatic residues involved (Bd4.3) or predicted to be involved (Bd5.1-7) in the interaction with FtsQ. The side chains of leucine residues (orange) that show important chemical shifts perturbations due to aromatic ring current effects of neighboring Trp or Tyr residues on the opposing β-strand are also displayed. β-sheet predictions from experimental data are highlighted in green and the yellow spheres indicate Hα protons for which interstrand NOEs (dotted black lines) were observed. For Bd4.3, an additional NOE between the amide group of the amidated C-terminal Pro residue and the acylated N-terminal Thr amide group is indicated. On the bottom right of each panel, the NMR solution structures of the peptides (light grey) are overlaid with (light blue) either the crystal structure (Bd4.3) or the AlphaFold 3.0 model (Bd5.1-7) in complex with FtsQ. High structural similarities with their corresponding structures/models are observed for both peptides. Propensities and order parameters were derived from backbone and CB chemical shifts using TALOS-N and RCI, respectively. The solution structures of Bd4.3 and Bd5.1-7 were calculated using CS-Rosetta from backbone and CB chemical shifts and interstrand distance constraints derived from experimental H***α***–H***α*** and H***α***–HN NOEs characteristic of antiparallel ***β***-sheets.

The structures in solution of both peptides were calculated with CS-Rosetta, using backbone and CB chemical shifts complemented with a few interstrand distance constraints derived for Hα-Hα and Hα-HN dipolar interactions (NOEs) of nuclei close in space (< 5 Å). The best 10 structures of each peptide are highly convergent, with an α-carbon atom mean pairwise RMSD ± STDEV of 0.13 ± 0.09 Å for Bd4.3 and of 0.88 ± 0.8 Å for Bd5.1-7. The best-energy structures are shown in (Figure 8). Both peptides adopt an antiparallel β-hairpin structure. In addition to backbone interstrand NOEs, several interstrand NOEs between sidechains and other NMR data support the calculated structures. For instance, for both peptides, leucine residue sidechains (highlighted in Figure 8 bottom panels) show highly perturbed chemical shifts due to ring current effects of the aromatic rings of nearby Tyr and Trp residues located in the opposite strand. For Bd4.3, a NOE is observed between the amidated C-terminal P19 residue and the amide proton of the acylated N-terminal Thr residue, demonstrating that both extremities of the peptide are close together.

In solution, Bd4.3 adopts effectively the same structure than in complex with FtsQ (Figure 8 bottom panel) with an α−carbon RMSD of 1.2 Å. Likewise, the solution structure of Bd5.1-7 resembles the AlphaFold 3.0 predicted structure in complex with FtsQ (RMSD of 0.58 Å). Importantly, for both peptides, the interstrand NOEs show that the structures of the peptides in solution and in the complex, whether experimental or predicted, are in frame and hence will expose similarly the sidechains for the interaction with FtsQ. Notably, NMR data indicate that RFdiffusion was harnessed to design peptides that adopt the predicted structure in solution and that are well poised to interact with FtsQ without major rearrangements or folding upon binding, which can be an asset for affinity avoiding entropic penalties, as well as for fast binding and competing with FtsL and FtsB.

### Design of Peptide Binders Targeting Pseudomonas aeruginosa FtsQ

To further assess the selectivity of the designed peptides for *E. coli* FtsQ, we examined their ability to interact with the orthologous *FtsQ* protein from *Pseudomonas aeruginosa* (FtsQ_*Pα*_; UniProt ID: G3XDA7). The DNA fragment encoding the periplasmic domain of FtsQ_*Pα*_ (residues 65–287) was cloned into pUT18C to generate pUT18C–FtsQ_*Pα*65-287_, and the resulting construct was co-transformed with pKT25-Bd5 Bd6, Bd7, Bd8 plasmids into *E. coli* DHT1 cells.

None of the Bd5–Bd8 peptides exhibited detectable interaction with the *P. aeruginosa* FtsQ periplasmic domain (FtsQ_*Pα*65-287_) in BACTH assays, demonstrating that these peptides are selective for *E. coli* FtsQ and do not associate with the *P. aeruginosa* orthologous protein (Figure 9A).

**Figure 9:**
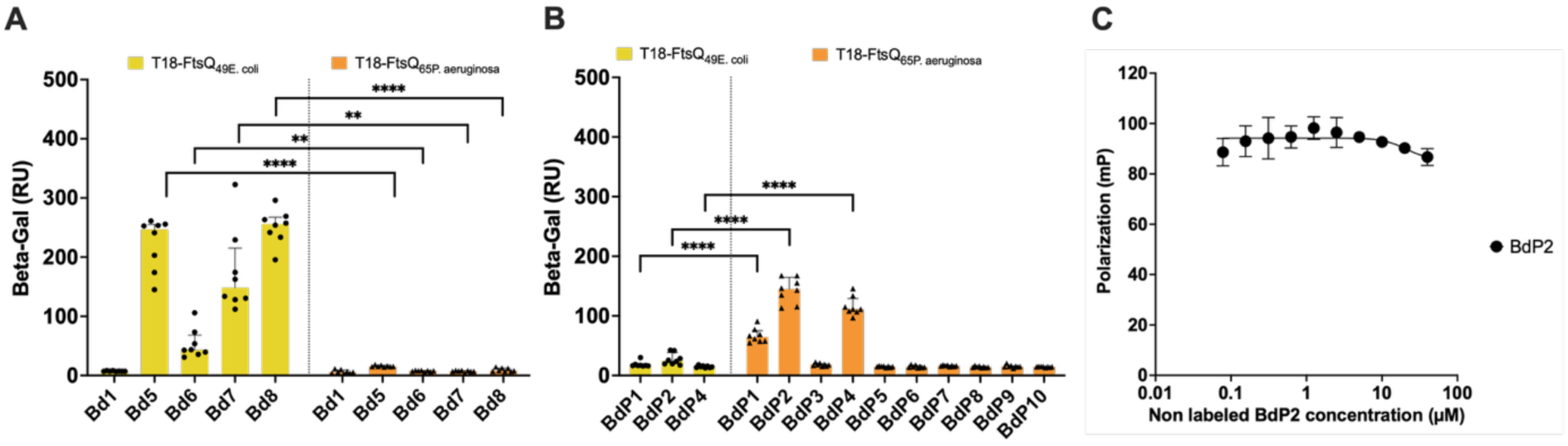
Analysis of peptides interaction according to bacteria species. **(A)** Bacterial two-hybrid (BACTH) analysis of interactions between the Bd peptides and either E. coli FtsQ (left) or Pseudomonas aeruginosa FtsQ (right). The β-galactosidase activities were measured on liquid cultures of E. coli DHT1 cells harboring the corresponding plasmids, as described in Materials and Methods. Each bar represents the median of 8 independent cultures (technical replicates), and error bars indicate the interquartile range. Brown-Forsythe and Welch One-way Anova test and Dunett T3 post hoc, ns > 0.05, **P < 0.01, ***P < 0.001 and ****P<0.0001. **(B)** BACTH analysis of interactions between P. aeruginosa–targeting peptides, BdP and the FtsQ proteins from either E. coli or P. aeruginosa. Same method than Fig7A **(C)** Fluorescence polarization (FP) competition assays evaluating the ability of increasing concentrations of unlabeled BdP2 (P. aeruginosa–targeting peptides) to compete with Cy5–Bd8 (50 nM) for binding to E. coli FtsQ_49-276_ (2 µM). Data points represent the mean of three independent measurements, with error bars indicating standard deviation. The data were fitted using a variable-slope dose–response inhibition model in Prism software. The curves depict one representative assay among three independent ones.

In parallel, we applied the same RFdiffusion pipeline to generate candidate peptide binders targeting FtsQ*_P.a_*. Ten sequences with the highest predicted scores were selected for experimental testing. Synthetic oligonucleotides encoding these peptides, designated BdP1 to BdP10, were cloned into the pKT25 vector, and the resulting constructs were tested in BACTH complementation assays with T18–FtsQ_Pa65-287_. Three peptides BdP1, BdP2, and BdP4 exhibited detectable interactions with the periplasmic domain of *P. aeruginosa* FtsQ, while showing no interaction with *E. coli* FtsQ (Figure 9B). Fluorescence polarization assays further confirmed that the synthetic BdP2 peptide did not bind the periplasmic domain of *E. coli* FtsQ *in vitro* (Figure 9C). Together, these results demonstrate that the RFdiffusion-based design strategy can be used to generate species-selective peptide binders targeting the essential cell division protein FtsQ.

## Discussion

The interaction between the C-terminal regions of FtsB (residues 64–87) and FtsQ (residues 246–272) plays a central role in divisome assembly and is essential for bacterial survival. This protein–protein interaction is mediated by β-strand augmentation between FtsB and FtsQ and is further stabilized by the β-strand contribution of FtsL. Therefore, disrupting this interface therefore represents an attractive strategy to interfere with bacterial cell division.

In this work, we exploited the RFdiffusion framework to design β-sheet peptides that conserve the key interaction hot spots of FtsB, with the goal of developing peptide-based inhibitors of the FtsQBL complex. Using a combination of in vitro biophysical assays (BLI and FP) and in vivo genetic interaction assays (BACTH), we demonstrated that several of the designed peptides, Bd4 to Bd8, bind selectively to the FtsB-binding site of FtsQ. Importantly, the BACTH interactions were detected with both the full-length FtsQ natively inserted in the inner membrane, and its isolated periplasmic domain expressed in cytosol. This indicates that Bd peptide–FtsQ binding does not rely on indirect membrane-associated interactions but reflects a direct molecular recognition event.

A notable exception was the peptide Bd4, which interacted with FtsQ only in the cytosolic BACTH assay. In this peptide, the FtsB-derived hot-spot motif is positioned toward the N terminus, in contrast to Bd5–Bd8, where it is located at the C terminus. This reversed orientation likely interferes with productive binding when Bd4 is tethered to a transmembrane segment for periplasmic targeting and highlights the importance of peptide topology and fusion geometry in functional assays. Further structural or biophysical analyses may help clarify the precise constraints imposed by peptide orientation in this context.

Mechanistically, our results demonstrate that the designed peptides bind specifically to the C-terminal region of FtsQ (residues 246–276). Alanine substitution of the aromatic residues F15 and Y16 in peptide Bd8—corresponding to the conserved F84 and Y85 anchors of FtsB—completely abolished FtsQ binding, both in vivo and in vitro. These findings indicate that the peptides engage the same interaction determinants as native FtsB. This conclusion was further supported by three-hybrid competition experiments, in which co-expression of both FtsB and FtsL fully disrupted the interaction between Bd8 and FtsQ, while FtsB alone was insufficient to do so. This observation is consistent with previous reports showing that the FtsQ–FtsB interaction is weak in the absence of FtsL and underscores the cooperative nature of the native FtsQBL assembly.

Crucially, the X-ray crystal structure of the Bd4.3–FtsQ complex provides direct structural validation of the binding mode generated by the RFdiffusion design strategy. The structure confirms that the peptide binds to the C-terminal β-strand interface of FtsQ and faithfully reproduces the β-strand augmentation geometry observed in the native FtsQBL complex. Beyond simple mimicry of FtsB, the designed peptide occupies a composite interaction surface spanning regions normally engaged by both FtsB and FtsL. In particular, the conserved tryptophan residue systematically introduced during the design process inserts into a hydrophobic pocket adjacent to the native interface and that is not exploited by the natural partners. This illustrates how generative design can combine interface mimicry with additional stabilizing interactions.

Complementary NMR analyses further demonstrate that peptides Bd4.3 and Bd5.1-7 adopt well-defined antiparallel β-hairpin conformations in solution. Importantly, the solution structure of Bd4.3 closely matches the conformation observed in the crystal structure of the FtsQ-bound complex, indicating that binding occurs without major folding rearrangement. Likewise, the solution structure of Bd5.1-7 shows strong agreement with the AlphaFold-predicted complex model. Together, these results indicate that RFdiffusion generated peptides that are structurally pre-organized prior to binding and geometrically compatible with the FtsQ interface, thereby reducing entropic penalties associated with folding upon binding and facilitating competition with the native FtsB/FtsL partners.

More generally, our results suggest that β-strand augmentation interfaces may represent particularly tractable targets for generative peptide design. In such interfaces, binding specificity emerges from conserved backbone hydrogen-bonding geometry combined with a limited number of hotspot side-chain interactions. This structural organization provides strong geometric constraints that can be efficiently captured during diffusion-based design while still allowing diversification of peptide topology and sequence context. The convergence of independently generated peptide architectures toward a common binding geometry, together with their validation by X-ray crystallography and NMR, supports the idea that these interfaces define transferable interaction frameworks that can be exploited to engineer competitive binders against multiprotein assemblies.

Importantly, successful binder generation required explicit incorporation of structural knowledge of the FtsQBL interface. Functional hotspot residues defining the β-augmentation register were first identified through literature survey of mutation studies interface analysis and ligandability/interactability mapping using our tool InDeep^23,24^. Initial attempts using unconstrained de novo RFdiffusion sampling did not yield peptides capable of reproducibly engaging the FtsQ binding groove. In contrast, constraining diffusion around the TFYRL β-strand hotspot motif derived from the native FtsB/FtsL interaction enabled the generation of multiple productive scaffolds that preserved the β-augmentation geometry while diversifying peptide topology. These results emphasize that generative approaches benefit strongly from interpretable structural priors and illustrate how predictive interface analysis and diffusion-based design can operate synergistically in protein–protein interaction targeting.

Recent studies have demonstrated the ability of diffusion-based approaches to generate high-affinity protein binders against folded protein surfaces. In contrast, the present work focuses on targeting a cooperative β-augmentation protein–protein interface within an essential bacterial multiprotein assembly and combines generative design with atomic-resolution structural validation and cellular functional assays. This integrated validation across structural and biological scales highlights the potential of constrained generative approaches for modulating dynamic interaction interfaces rather than isolated binding surfaces.

Importantly, this binding mode reflects a cumulative mimicry mechanism in which a single designed peptide substitutes for structural contributions normally provided by two distinct proteins, FtsB and FtsL. While conserved aromatic hotspot residues reproduce the anchoring interactions of FtsB, the surrounding scaffold generated by RFdiffusion extends across the composite interface and partially recapitulates stabilizing contacts associated with FtsL. Rather than competing with a single interaction partner, the designed peptides effectively engage the cooperative interaction surface underlying FtsQBL assembly. This observation emphasizes how generative design approaches can exploit composite protein–protein interfaces to produce binders that integrate multiple interaction determinants within a single molecular scaffold.

The initial set of linear peptides (Bd4, Bd5, and Bd8) bound FtsQ with dissociation constants ranging from the 0.4 to 1.4 µM range, representing a substantial improvement over the affinity reported for the linear FtsB(64–87) peptide (Kd ≈ 9 µM) ^16^. However, despite these affinities, the first-generation peptides were unable to disrupt divisome assembly or induce a detectable phenotype when overexpressed in bacteria. This limitation is not unexpected, as even a macrocyclic FtsB-derived peptide with submicromolar affinity has been shown to lack antibacterial activity^16^. These observations underline that binding affinity alone is insufficient to achieve functional inhibition in the cellular context.

To overcome this limitation, we performed a second round of RFdiffusion-based optimization, leading to peptides with significantly improved functional activity. Optimized peptides such as Bd5.1 and Bd8.3 exhibited enhanced interaction signals in BACTH assays, reduced IC_50_ values in FP experiments, and induced pronounced filamentation and growth defects when overexpressed in bacteria. These phenotypes are characteristic of impaired cell division and are consistent with effective disruption of FtsQBL assembly.

We further enhanced the antimicrobial activity of the peptides introducing positively charged residues to facilitate entry into the bacterial periplasm. This strategy proved effective: the optimized peptide Bd5.1-7 fully inhibited the growth of the permeabilized *E. coli* lptD4213 strain at micromolar concentrations and induced a characteristic chaining phenotype indicative of division arrest. Importantly, alanine-substituted and scrambled control peptides lacked both antibacterial activity and FtsQ binding, demonstrating that toxicity is driven by specific target engagement rather than nonspecific effects, such as membrane lysis. Moreover, Bd5.1-7 showed no detectable cytotoxicity toward human lung fibroblasts, supporting its selectivity.

At present, we do not know the fraction of peptide that reaches the periplasm, nor its stability once inside the bacterial envelope. These parameters will be critical for further development and clearly motivate the design of more stable peptidomimetic derivatives resistant to proteolytic degradation. Nonetheless, our results establish a clear proof of concept that rationally designed peptides can access and functionally inhibit a periplasmic divisome target in Gram-negative bacteria.

Finally, by leveraging predictive interface mapping using InDeep ^23,24^ to identify transferable hotspot determinants within the FtsQ β-augmentation interface, together with RFdiffusion-based scaffold generation, we demonstrated that it is possible to generate peptides that selectively target FtsQ orthologs from *E. coli* or *Pseudomonas aeruginosa*. This species selectivity opens the possibility of developing narrow-spectrum antibacterial agents that spare commensal microbiota and reduce selective pressure for resistance. More broadly, this work illustrates how interpretable interface analysis coupled to AI-driven generative design and experimental validation can enable the rational targeting of challenging protein–protein interactions in bacteria.

## Limitations

Despite the encouraging biological and structural results obtained in this study, several aspects remain to be clarified before further translational development. In particular, the fraction of externally added peptide that effectively reaches the bacterial periplasm remains unknown, as does its stability within the bacterial envelope. These parameters are likely to influence the apparent cellular potency observed in growth inhibition assays. Finally, while antimicrobial activity was demonstrated in a permeabilized *E. coli* strain, further work will be required to evaluate peptide uptake, stability, and activity in more physiological bacterial contexts.

## Perspectives

The results presented here establish a framework for the rational targeting of divisome protein–protein interactions using AI-guided peptide design. Future work will focus on improving peptide pharmacological properties through the development of stabilized peptidomimetics and constrained scaffolds aimed at enhancing resistance to proteolysis and bacterial uptake. More broadly, this work illustrates how AI-driven generative design can be integrated within a complete experimental validation framework spanning structural prediction, atomic-resolution characterization, and functional perturbation of a native multiprotein assembly. By combining hotspot-guided design, structural validation by X-ray crystallography and NMR, quantitative biophysics, and cellular phenotyping, our results establish a closed design-to-function pipeline for targeting protein–protein interfaces in bacteria. Such integrated strategies may help transform structurally characterized but pharmacologically underexploited multiprotein assemblies into tractable targets for antibacterial development.

## Material and Methods

### RF-diffusion-based design of FtsB/L mimetic peptides targeting FtsQ

The periplasmic domain of *Escherichia coli* FtsQ was used as the fixed receptor for peptide design. The surface groove on FtsQ that accommodates the native FtsB/L β-strand was defined as the design target. Using InDeep, we first characterized the interaction fingerprint of the native FtsB/L–FtsQ interface, identifying the β-strand segment TFYRL as a minimal recognition element. This motif was then extracted and used as the seed for the motif-scaffolding designs described below.

To generate peptide backbones two design scenarios were explored. In Series 1, native-like mimics were generated by modeling conservative substitutions around the native FtsB/L β-strand, in order to assess how closely the native geometry could be retained while remaining synthetically tractable. In Series 2, a motif-scaffolded approach was used in which the TFYRL β-strand motif was pre-positioned in the FtsQ groove in a strand-augmentation geometry, and RFdiffusion was instructed to generate the flanking backbone segments around this locked motif. Around 1,000 peptide backbones were generated. Each backbone was subsequently assigned two sequences using ProteinMPNN in complex mode, keeping the FtsQ receptor fixed. The binder-to-receptor contact mask was defined to include the FtsQ groove region to promote productive strand extension and complementarity at the interface.

All designed peptide–FtsQ complexes were rapidly evaluated using AlphaFold2. Complexes were retained if the interface predicted alignment error (pAE) was below 10 (median across contacting residues), if no severe steric clashes were observed upon visual inspection, and if the designed peptide formed a clear β-strand augmentation into the FtsQ groove. Surviving models were clustered by backbone RMSD and sequence identity to ensure structural and sequence diversity, and representative designs were selected from each cluster for further consideration.

After filtering and clustering, 8 peptides were selected across the two design tracks. Series 1 included the native-like FtsB/L mimics and Series 2 contained the motif-scaffolded designs incorporating the locked TFYRL strand with diversified flanking regions.

### Bacterial strains and growth media

The *E. coli* K-12 strain XL1-Blue (Stratagene) was used in all of the cloning steps. BACTH complementation assays were carried out either with the *E. coli* cya strain DHM1 [F−glnV44(AS) recA1 endA gyrA96 thi-1 hsdR17 spoT1 rfbD1 cya-854] mostly for BACTH with full-length FtsQ or with the DHT1 [F-, cya-854, ilv 691::Tn10, recA1, endA1, gyrA96(Nal r), thi1,hsdR17, spoT1, rfbD1, glnV44(AS)] for BACTH experiments with FtsQ periplasmic domain.

Bacteria were grown at 30°C with shaking at 180rpm in Luria-Bertani (LB) broth supplemented with ampicillin at 100 μg/ml and kanamycin at 50 μg/ml as needed. For cloning steps, glucose (0.2%) was added to the growth medium to decrease the expression of hybrid proteins. Screening for β-galactosidase expression was performed on LB agar plates supplemented with 5-bromo-4-chloro-3-indolyl-β-d-galactoside (X-Gal; 40 μg/ml), isopropyl-β-d-galactopyranoside (IPTG; 0.5 mM), and appropriate antibiotics.

For protein production, the *E. coli* K-12 strain Krx strain was used. This strain contains a chromosomal copy of the T7 RNA polymerase gene tightly regulated by a rhamnose promote (rhaPBAD) while the proteins critical for rhamnose metabolism [isomerase (RhaA), kinase (RhaB) and aldolase (RhaD)] are deleted. Bacteria were grown at 30°C with shaking at 180rpm in Luria-Bertani (LB) broth supplemented with kanamycin at 50 μg/ml and required concentration of glucose and rhamnose.

For bacteria morphological analysis and growth monitoring when peptides are overproduced in the cell, the *E. coli MG1655* [F− ilvG rfb-50 rph-1] strain was used.

To test the efficiency of the synthetic peptides the *E. coli K12-* NR698 [MC4100 *lptD4213*] strain was used as it exhibits an increased outer membrane permeability^22^.

### Plasmid constructions

pKT25-ZIP, pKTM25-ZIP pUT18C-FtsQ_1-276_ and pUT18C-FtsQ_1-246_ were constructed in prior studies as indicated in Table S5.

For the constructions of the other plasmids standard protocols for molecular cloning, PCR, DNA analysis, and transformation were used. PCR was performed with Phusion™ Hot Start II DNA Polymerase (Thermo Fischer Scientific). Oligonucleotides were from Sigma Aldrich and synthetic genes from TwistBioScience. All the DNA sequences encoding for the designed peptides (Bd1 to Bd8) have been synthesized as single-stranded oligonucleotides (forward and reverse) by Sigma Aldrich and annealed to form double-stranded DNA by our team (TableS4)

The plasmids pKTM25-X and pKT25-X (with X corresponding to the DNA sequences encoding for peptide Bd1 to Bd8) were constructed by cloning the oligonucleotides between the *BamH*I and the *Acc65*I restriction sites of the pKTM25 and pKT25 vectors. For the plasmid pUTM18C_Bd5/8/51-7, the oligonucleotide sequence was inserted into between the *BamH*I and the *EcoR*I cloning sites of the pUTM18C_ZIP vector. For the plasmid pUTM18C_Bd4.3/5.1/8.3, the DNA sequences encoding the peptides were amplified by PCR from the corresponding pKT25-X plasmids using optimized peptides primers (Table S6).

To construct the plasmids, pUT18C_FtsQ*_49-276_* and pUT18C_FtsQ*_49-246,_* the DNA sequences encoding the FtsQ_49-276,_ and FtsQ*49-246* truncated fragments were amplified by PCR from *E. coli* genome using appropriate primers containing XbaI and EcoRI restriction sites, and inserted into pUT18C vector linearized by the same restriction enzymes. The plasmid pUT18C-Fts*Q_P.a65-285_* encoding for the hybrid T18-FtsQ*_P.a_*_65-285_ cytosolic mutant of FtsQ from *P.aeruginosa* was constructed similarly, except that BamHI and Acc65I restriction sites were used. The plasmid pKTN*-*Fts*Q* was built by cloning the wild type *ftsqQ* gene into pKNT25 vectors using PstI and Bsu15I restriction enzymes. For pETM-FTsQ_49-276_, *ftsQ* was amplified by PCR and cloned using the Gibson assembly method into pETM-11 cut by the *Nco*I and *Acc65*I restriction enzymes^25^.

The plasmid pKTM25-Bd8-T18-FtsQ, that co-expresses the two hybrid partners (T25-TM-Bd8 and T18-FtsQ), was created by Gibson assembly using PCR amplified *T18-ftsQ* gene with primers containing homology sequence to the vector pKTM25_Bd8 digested by EcoRI and PvuI.

The plasmids pUC19-FtsB, pUC19-FtsL and pUC19-FtsB-SD-FtsL, expressing respectively FtsB, FtsL or both proteins, were generated by inserting a short double stranded oligonucleotide sequence (pUC19oligo, Table S6) between the *Hind*III and *EcoR*I restriction sites of plasmids pUT18C-FtsB, pUT18C-FtsL or pUT18C-FtsB_SD-FtsL, thus deleting the T18 coding fragment (Table S5).

To construc the 30 optimized peptides deriving from Bd4 (Bd4.1 to Bd4.10), Bd5 (Bd5.1 to Bd5.10) and Bd8 (Bd8.1 to 8.10), 10 DNA sequences (Table S7), each one encoding for three of the optimized peptides (one in frame, the two other in 3’extremity out of frame) were ordered and cloned into pKT25 plasmid between the *Pst*I and *EcoR*I restriction sites. By digesting with NheI and religating the plasmids, the second encoding sequence were put in frame while the plasmids were digested by both NheI and SpeI to put in frame the third sequence. A similar process was used for the peptides targeting *P.aeruginosa* FtsQ or Bd8 mutants. To clone the optimized peptides into pKTM25 vectors, dna sequences were amplified by PCR using appropriate primers and fused using Gibson assembly into pKTM25 cut between *Xba*I and *EcoR*I resitrictions sites.

Each plasmid’s DNA sequence, once constructed, was confirmed by sequencing (Eurofins Genomics, Cologne Germany).

### BACTH complementation assays

For BACTH complementation assays, recombinant pKT25 (or pKTM25) expressing the T25-Bd peptides or T25-TM-Bd peptides and pUT18C derivatives expressing the T18-FtsQ or T18-FtsQ_49-246,_ fusions were co-transformed by electroporation into DHM1/ DHT1 cells. The transformants were plated on LB plus X-Gal (40μg/mL) and IPTG (0.5mM) with ampicillin and kanamycin and incubated at 30°C. To identify the presence of an interaction between the FtsQ protein and the designed peptides, the functional complementation between the hybrid proteins was quantified by measuring ß-galactosidase activities in bacterial liquid culture^26^. These measurements were carried out in 2.2 mL 96- deep-well plate. Eight independent colonies from each set of transformations were picked and inoculated in 300μL sterile LB broth supplemented with 0.5 mM IPTG, kanamycin and ampicillin. The 96-wells plates were sealed with a microporous tape sheet to allow gas exchange and incubated overnight at 30°C with shaking at180 rpm.

The next day, the cultures were diluted fivefold by adding 1.2 mL of M63 medium in each well and the optical density at 600 nm (OD600) was recorded. In parallel, 7μL of 0.05% SDS + 10μL of chloroform were added into 200 μL of the diluted bacterial suspensions to permeabilize the cells. The mixture was mixed 20 times by pipetting vigorously and left under a fume hood at RT for one hour to allow chloroform evaporation. For the enzymatic reaction, 105 μL of PM2 reaction buffer (containing 100 mM β-mercaptoethanol, and 0.1% o-nitrophenol-β-galactoside (ONPG)) were mixed with 20 μL of the permeabilized cells. After 10-20 minutes incubation at RT (until sufficient yellow color has developed), the reaction was stopped by adding 50 μL 1 M Na2CO3 and the absorbance data at 405 nm (OD405) with the microplate reader FLUOstar Omega BMG LABTECH. To quantify β-galactosidase activity.

To calculate the level of β-galactosidase activity, the following formula was used: ß-gal = (OD_405nm_)x1000 / (OD_600nm_)xt_(min)_

### Protein production and purification

For protein production, the plasmid pETM-FtsQ_49_ plasmid was construced to express the FtsQ_49_ mutant fused at its N-terminal extremity to a 6xHis Tag and a a Tobacco Etch Virus (TEV) protease cleavage site. The PCR amplified *ftsQ_49_* fragment was cloned by using Gibson assembly into the pETM-11 vector digested by *Nco*I and *Acc65*I.

The *E. coli KrX* strain was then transformed with the pETM-FtsQ_49_ plasmid and grown OVERNIGHT in LB medium supplemented with kanamycin and glucose (0.2%) at 30°C with shaking at 180rpm. To induce the protein expression, the OVERNIGHT culture was diluted 100 times and grown in 1 liter of LB media supplemented with kanamycin, 0.05% glucose and 0.1% of rhamnose for 7 hours at 30°C with shaking at 180 rpm. Bacteria were collected by 20 min centrifugation at 10 000 rpm.

Bacterial cell were resuspended in 20 mL of buffer (20 mM Hepes, 150 mM NaCl, pH 7.5) supplemented with a protease inhibitors cocktail from Merck. After lysis by sonication, bacteria were centrifugated 20 minutes at 10,000 rpm at 4°C. The supernatant was loaded on a Ni-NTA column (Qiagen) equilibrated with 5mM Imidazole, 20 mM Hepes, 150 mM NaCl, 7.5 pH. The column was then washed with equilibration buffer. Proteins were eluted in 100mM Imidazole, 20 mM Hepes, 150 mM NaCl, 7.5 pH buffer supplemented with protease inhibitor. Collected fractions were analyzed by SDS-PAGE on NuPAGE^TM^ 4-12% Bis-Tris Gel (Invitrogen) (Figure S1). The fraction containing the purified protein were concentrated by ultracentrifugation (10kDa tubes Amicon Sigmal-Aldrich) for 1 hour at 3000rpm at 4°C. The concentrated solution was finally dialyzed against 20 mM Tris, 150 mM NaCl, pH8 buffer using 10,000MWCO dialysis cassette G2 (ThermoScientific). The concentrated purified proteins were stored at 4°C.

For X-ray crystallography experiment, the 6xHis tag was then cleaved oi by the TEV protease digestion (ratio 1:100 (w/w) in the presence of 1mM DTT. The mixture was incubated at 11° С OVERNIGHT. To eliminate the cleaved tag and the TEV protease, the digested mixture was passed over to a second Ni-NTA column and the flow-through was collected.

### BioLayer Interferometry

BioLayer Interferometry (BLI) measurements were performed on an Octet HTX (Sartorius) at room temperature, with orbital shaking at 1,000 rpm. Biotinylated peptides (Bd1, Bd4, Bd5 and Bd8) were synthetized by Genosphere Biotechnologies (Paris, France).

Two strategies were employed: Immobilization of peptides onto Streptavidin coated biosensor (SA) or immobilization of the FtsQ_49_ protein onto Ni-NTA biosensor (Ni-NTA biosensors, Satorius).

Biosensors were equilibrated in running buffer for at least 10 min prior to use (20mM Tris-HCl, 150mM NaCl, 0.1% BSA, 0.01% polysorbate 20 (P20), pH8.0), which was also used throughout the experiments. For peptides immobilization, peptides were loaded at required concentration (between 30 and 67ng/mL) for 300 seconds to reach a loading response of 0.5-1 nm, followed by a 300s equilibration in running buffer. Association of 6His-FtsQ_49–276_ to immobilized peptides was recorded by dipping sensors into a 1:3 serial dilution series starting at 5 µM 6His-FtsQ_49–276_ for 1,200 s, followed by dissociation in running buffer for 1,200 s. Signals were corrected by subtracting the response from reference sensors and buffer-only wells (double referencing). Global fitting across all analyte concentrations was performed in Octet Analysis Studio using a 2:1 heterogeneous ligand model.

### Fluorescence Polarization assay

FP measurements were performed in FP buffer (20mM Tris-HCl, 150mM NaCl, 0.1%, BSA, 0.01% polysorbate 20 (P20), pH8.0). Cy5-labeled peptides were diluted in FP buffer to a final concentration of 50 nM in 30 µl of total volume reaction per well and dispensed a black, flat bottom, non-binding surface 384-well plate (Corning). Twofold serial dilution series of FtsQ_49–276_, prepared starting at 20 µM in FP buffer, were added to the peptide-containing wells. Fluorescence polarization was recorded on a Tecan Spark multimode plate reader equipped with polarizing filters using at 635 nm and emission at 665 nm. Apparent dissociation constant (K_D_) was determined by nonlinear regression analysis of one-site total binding curves in GraphPad Prism software.

For the competition experiments, untagged peptides or biotinylated peptides (Bd1, Bd4, Bd5 and Bd8; non fluorescent, previously used for BLI) were titrated against Cy5-labeled peptide (Bd8) in the presence of 6His-FtsQ_49-276_. In 384-well plates as above, 10 µl of Cy5-Bd5 solution (final concentration 50nM), 10 µl of 6His-FtsQ_49-276_ solution (final concentration 0.5µM) and 10 µl of a two-fold serial dilution of competitor peptides were mixed in a final volume of 30 µl per well. Fluorescence polarization was measured using the same optical settings as above. Half-maximal inhibitory concentrations (IC_50_) were determined by nonlinear regression using the [inhibitor] vs response (variable slope, 4-parameter) model in GraphPad Prism software.

### Phenotypic analysis

To test the effects of Bd peptide overproduction in bacteria, chemically competent *E. coli* MG1655 cells were transformed with pUTM18C_BdX plasmids. For each transformant, several clones were collected and cultivated OVERNIGHT in 5mL of LB + ampicillin (100µg/mL) + glucose 0.2% at 30°C, 180rpm. The following morning, the pre-culture was diluted 1:100 in 5mL of LB + ampicillin (100µg/mL) and after 30 min of growth at 30°C 180 rpm, IPTG was added (0.5mM final concentration) to induce the expression of the BdX fusion proteins. The bacteria were observed by phase contrast microcopy after 3h30 and 6h30 of culture. For this, 1 ml of bacterial cultures were briefly centrifuged and the cell pellet was resuspended in MB631X media and 2µL deposited on a glass slide and covered by coverslip. Bacteria were imaged with a Nikon epi-fluorescence microscope Eclipse 80i equipped with a 100x Plan-Apo oil immersion objective. The captured images were subsequently analyzed with Fiji and the ObjectJ Cell Counter plugin. For the bacteria with a wildtype phenotype (around 4µm and below) the automatic Coli-Inspector macro was used. For filamentous bacteria, the length was manually measured^27^.

The effects of synthetic Bd peptides on cell division and bacterial growth were tested on the *E. coli* imp4213 strain that exhibits increased outer-membrane permeability^22^. The *E. coli* imp4213 cells were grown overnight in 5 ml of LB at 37°C and 180 rpm. The day after the overnight culture was diluted 1:1000 in 5mL LB and cultivated for 2 hours at 37°C and 180 rpm. Aliquots of 200µL of bacterial culture were transferred into 2mL Eppendorf tubes and the Bd peptides were added at a concentration equal to 2 times their MIC. A negative control was performed using DMSO at similar concentration. The tubes were cultured at 37°C at 200 rpm. At each time point the corresponding tubes were collected and bacteria harvested for microscopic analysis as described above.

### Minimal Inhibitory Concentration Measurement

*E. coli* imp4213 cells were grown overnight in 5 ml of LB at 37°C and 180 rpm. The day after the overnight culture was diluted 1:1000 in 5mL LB and cultivated for 2 hours at 37°C and 180 rpm. The Bd peptides to be tested were diluted (from a 10 mM stock solution in DMSO) in LB to a final concentration of 200 µM. Then, 2-fold serial dilutions with 8 points starting from 50µM were prepared in 100µL of LB in 96-well plates with flat bottom. The bacterial pre-culture was inoculated at a 1:100 dilution in each well, and culture were grown at 37°C and 200 rpm for 16 hrs in an Omega Fluostar microplate reader with measurement of OD_600nm_ every 15 minutes. DMSO (at equivalent concentration) was used as negative control.

### Cytotoxicity Assay

Cytotoxicity was determined using the CellTiter-Glo luminescent cell viability assay (Promega). White with clear bottom 384-well plates were seeded with 2×10^3^ MRC5 human normal lung fibroblast cell per well in a final volume of 20µL. The next day, the individual compounds were dispensed into 384-well plate using the Echo 550 acoustic dispenser at concentrations 2X. DMSO-only (0.5%) and camptothecin (25µM; Sigma-Aldrich) controls were added. The compounds were resuspended in 40µL/well of culture media and 20µL were transferred in the 384-well plates containing the MRC5 cells. After 48h incubation, 20 µl/well of Celltiter-Glo reagent was added, incubated for 20 min and luminescence was recorded using a Mitras luminometer (Berthold Technologies) with 0.5 sec.

### Protein crystallization

Crystallization screening was performed by the vapor-diffusion method using a Mosquito nanodispensing system (SPT Labtech) following established protocols^28^. Initial hits were optimized by hanging-drop vapor diffusion at 18 °C. The best crystals were obtained by mixing the protein complex with reservoir solution containing 10%(w/v) PEG20K, 0.1M Bicine pH9, and 2% (v/v) dioxane. Crystals were cryoprotected with reservoir solution supplemented with 33% (v/v) ethylene glycol and flash-cooled in liquid nitrogen.

### X-ray data collection and structure determination

X-ray diffraction data were collected at beamlines PROXIMA 1 and PROXIMA 2A (Synchrotron SOLEIL, Saint-Aubin, France) and processed with autoPROC^29^. The structure was solved by molecular replacement using Phaser^30^ with an AlphaFold3 model as the search template. The final model was built through iterative cycles of manual rebuilding in Coot^31^ and reciprocal-space refinement with BUSTER^32^. Atomic coordinates and structure factors have been deposited in the Protein Data Bank under accession codes.

### NMR

NMR experiments were recorded on a 600 MHz Avance NEO (Bruker Biospin, Billerica, USA) spectrometer with a 14.1 Tesla magnetic field. The spectrometer was equipped with a cryogenically cooled triple resonance ^1^H[^13^C/^15^N] probe. Spectra were recorded and processed using TopSpin 4.5 (Bruker Biospin) and analyzed using CCPNMR 3.3.3^33^. Experiments were performed at 25°C. The peptides (Genosphere) were acetylated on the N-termini and amidated on the C-termini. Peptide Bd4.3 (517 µM) was dissolved in 20 mM sodium phosphate pH 6.2, 3% D_2_O (Eurisotop) and peptide Bd5.1-7 (667 µM) in 10 mM sodium phosphate, 20 mM deuterated Tris-HCl (Tris-d18, Eurisotop) pH 6.3, 150 mM NaCl, 3% D_2_O. For assignment, structural characterization and determination, homonuclear ^1^H-^1^H TOCSY (Total Correlation SpectroscopY, 80 ms mixing time) and NOESY (Nuclear Overhauser Effect SpectroscopY, 250 ms mixing time), and natural abundance heteronuclear ^1^H–^15^N (SOFAST) and ^1^H–^13^C (Edited-HSQC) correlation spectra were acquired using standard pulse sequences from the Bruker and NMRLib^34^ libraries. Per-residue secondary structure propensities and S^2^ order parameters reflecting backbone rigidity/flexibility were derived from backbone and CB chemical shifts using TALOS-N^35^ and RCI^36^, respectively. Solution structures were calculated with CS-Rosetta^37^ from backbone chemical shifts and sparse backbone distance constraints derived from experimental NOEs. The best 10 structures were chosen to describe the structure of the peptides. Structures were visualized with ChimeraX^38^ and graphs were plotted with GraphPad Prism 10.6.0.

### Statistical analysis

For β-galactosidase activity and bacteria length measurements, the results are presented as median ± interquartile. For FP and cytotoxicity experiment, all data are presented as the means ± standard error. Statistical significances were analyzed using a one-way analysis of variance (ANOVA) with Brown-Forsythe and Welch correction, and post hoc Bonferroni test Student’s t test. For multiple comparison with sample n<50 (BACTH assay) Dunnett T3 post hoc was used while multiple comparison with sample n>50 (morphological assay) Games-Howell post hoc was used. Statistics were evaluated with GraphPad Prism 10.6.1. A P value of <0.05 was considered statistically significant. Figures were illustrated using the same software.

## Acknowledgments

This research was funded by Institut Pasteur and the Centre National de la Recherche Scientifique, CNRS UMR 3528 (Biologie Structurale et Agents Infectieux) and Institut Pasteur. (grant number CACSICE Equipex ANR-11-EQPX- 0008).

This work benefited from a government grant (PPR Antibioresistance) managed by the Agence Nationale de la Recherche – Programme d’Investissements d’Avenir (ANR-20-PAMR-0011)

The funders had no role in study design, data collection and analysis, decision to publish, or preparation of the manuscript.

P.R. was supported by Université Paris Cité, F-75006 Paris

F.C. was supported by Institut Pasteur (grants PTR566).

The authors thank T. Silhavy for sharing the *E. coli lptD4213 strain*.

A CC-BY public copyright license has been applied by the authors to the present document and will be applied to all subsequent versions up to the Author Accepted Manuscript arising from this submission, in accordance with the grant’s open access conditions. »

## Author contributions

Conceptualization: O.S., D.L., X.L., P.R. and F.C. Methodology: P.R., X.L., G.K., J.C., D.L. and O.S. Investigation: P.R., X.L., F.C., A.M., M.H.N., M.D., G.K., C. B. C., A.B., F. A., C.G., A.H. and J.C., Software: X.L. Visualization: P.R., O.S., and A.M. Supervision: D.L., O.S., J.C., A.H., J.I.G., and F.A. Writing—original draft: P.R., O.S., M.N., A.H. Writing—review and editing: D.L., O.S., G.K.,

## Data and material availability

All data supporting the conclusions of this study are provided in the article and the Supporting Information. Raw data generated during the study are available from the corresponding authors upon reasonable request

## References

1. Hutchings, M., Truman, A. & Wilkinson, B. Antibiotics: past, present and future. Curr. Opin. Microbiol. 51, 72–80 (2019).

2. Murray, C. J. et al. Global burden of bacterial antimicrobial resistance in 2019: a systematic analysis. The Lancet 399, 629–655 (2022).

3. Buckley, G. J. & Palmer, G. H. COMBATING ANTIMICROBIAL RESISTANCE AND PROTECTING THE MIRACLE OF MODERN MEDICINE A Consensus Study Report of. 10.17226/26350 (2022) doi:10.17226/26350.

4. World Health Organization. WHO Bacterial Priority Pathogens List, 2024- Bacterial pathogens of public health importance to guide research, development and strategies to prevent and control antimicrobial resistance. World Health Organization 1–72 (2024).

5. Attaibi, M. & Den Blaauwen, T. An Updated Model of the Divisome: Regulation of the Septal Peptidoglycan Synthesis Machinery by the Divisome. International Journal of Molecular Sciences 2022, Vol. 23, Page 3537 23, 3537 (2022).

6. Karimova, G., Dautin, N. & Ladant, D. Interaction network among Escherichia coli membrane proteins involved in cell division as revealed by bacterial two-hybrid analysis. J. Bacteriol. 187, 2233–2243 (2005).

7. Carro, L. Recent Progress in the Development of Small-Molecule FtsZ Inhibitors as Chemical Tools for the Development of Novel Antibiotics. Antibiotics 8, 217 (2019).

8. Käshammer, L. et al. Cryo-EM structure of the bacterial divisome core complex and antibiotic target FtsWIQBL. Nature Microbiology 2023 8:6 8, 1149–1159 (2023).

9. Käshammer, L. et al. Cryo-EM structure of the bacterial divisome core complex and antibiotic target FtsWIQBL. Nature Microbiology 2023 8:6 8, 1149–1159 (2023).

10. Nguyen, H. T. V. et al. Structure of the heterotrimeric membrane protein complex FtsB-FtsL-FtsQ of the bacterial divisome. Nature Communications 2023 14:1 14, 1903- (2023).

11. Kureisaite-Ciziene, D. et al. Structural analysis of the interaction between the bacterial cell division proteins FTSQ and FTSB. mBio 9, (2018).

12. Choi, Y. et al. Structural Insights into the FtsQ/FtsB/FtsL Complex, a Key Component of the Divisome. Scientific Reports 2018 8:1 8, 18061- (2018).

13. Bart van den Berg van Saparoea, H., et al. Fine-mapping the contact sites of the Escherichia coli cell division proteins FtsB and FtsL on the FtsQ protein. Journal of Biological Chemistry 288, 24340–24350 (2013).

14. Buddelmeijer, N. & Beckwith, J. A complex of the Escherichia coli cell division proteins FtsL, FtsB and FtsQ forms independently of its localization to the septal region. Mol. Microbiol. 52, 1315–1327 (2004).

15. Muñoz, K. A. et al. A Gram-negative-selective antibiotic that spares the gut microbiome. Nature 2024 630:8016 630, 429–436 (2024).

16. Paulussen, F. M. et al. Covalent Proteomimetic Inhibitor of the Bacterial FtsQB Divisome Complex. J. Am. Chem. Soc. 144, 15303–15313 (2022).

17. Bourbon, L. M. L., Villoutreix, B. O. & Sperandio, O. Which Three-Dimensional Characteristics Make E ffi cient Inhibitors of Protein − Protein Interactions? (2014).

18. Sperandio, O., Reynès, C. H., Camproux, A. C. & Villoutreix, B. O. Rationalizing the chemical space of protein-protein interaction inhibitors. Drug Discov. Today 15, 220–229 (2010).

19. Reynès, C. et al. Designing focused chemical libraries enriched in protein-protein interaction inhibitors using machine-learning methods. PLoS Comput. Biol. 6, (2010).

20. Watson, J. L. et al. De novo design of protein structure and function with RFdiffusion. Nature 2023 620:7976 620, 1089–1100 (2023).

21. Kureisaite-Ciziene, D. et al. Structural analysis of the interaction between the bacterial cell division proteins FTSQ and FTSB. mBio 9, (2018).

22. Ruiz, N., Falcone, B., Kahne, D. & Silhavy, T. J. Chemical Conditionality: A GeneticStrategy to Probe Organelle Assembly. Cell 121, 307–317 (2005).

23. Mallet, V. et al. InDeep: 3D fully convolutional neural networks to assist in silico drug design on protein–protein interactions. Bioinformatics 38, 1261–1268 (2022).

24. Mareuil, F. et al. InDeepNet: a web platform for predicting functional binding sites in proteins using InDeep. Nucleic Acids Res. 53, W324–W329 (2025).

25. Gibson, D. G. et al. Enzymatic assembly of DNA molecules up to several hundred kilobases. Nature Methods 2009 6:5 6, 343–345 (2009).

26. Karimova, G., Pidoux, J., Ullmann, A. & Ladant, D. A bacterial two-hybrid system based on a reconstituted signal transduction pathway. Proc. Natl. Acad. Sci. U. S. A. 95, 5752–5756 (1998).

27. ObjectJ Cell Counter. https://sils.fnwi.uva.nl/bcb/objectj/examples/CellCounter/cellcounter-md/cellcounter.html.

28. Weber, P. et al. High-Throughput Crystallization Pipeline at the Crystallography Core Facility of the Institut Pasteur. Molecules 24, (2019).

29. Vonrhein, C. et al. Data processing and analysis with the autoPROC toolbox. Acta Crystallogr. D Biol. Crystallogr. 67, 293–302 (2011).

30. McCoy, A. J. et al. Phaser crystallographic software. J. Appl. Crystallogr. 40, 658–674 (2007).

31. Emsley, P. & Cowtan, K. Coot: model-building tools for molecular graphics. Acta Crystallogr. D Biol. Crystallogr. 60, 2126–2132 (2004).

32. BUSTER version 2.10.0 – ScienceOpen. https://www.scienceopen.com/document?vid=34a668bc-6e6f-4572-a548-6d19e78e1e30.

33. Skinner, S. P. et al. CcpNmr AnalysisAssign: a flexible platform for integrated NMR analysis. Journal of Biomolecular NMR 2016 66:2 66, 111–124 (2016).

34. Favier, A. & Brutscher, B. NMRlib: user-friendly pulse sequence tools for Bruker NMR spectrometers. J. Biomol. NMR 73, 199–211 (2019).

35. Shen, Y. & Bax, A. Protein structural information derived from nmr chemical shift with the neural network program talos-n. Methods in Molecular Biology 1260, 17–32 (2015).

36. Berjanskii, M. V. & Wishart, D. S. A Simple Method To Predict Protein Flexibility Using Secondary Chemical Shifts. J. Am. Chem. Soc. 127, 14970–14971 (2005).

37. Shen, Y. et al. Consistent blind protein structure generation from NMR chemical shift data. Proc. Natl. Acad. Sci. U. S. A. 105, 4685–4690 (2008).

38. Pettersen, E. F. et al. UCSF ChimeraX: Structure visualization for researchers, educators, and developers. Protein Science 30, 70–82 (2021).

