## supplementary_file for "Structure-guided generative design of peptides targeting the FtsQBL divisome complex inhibit *Escherichia coli* cell division"

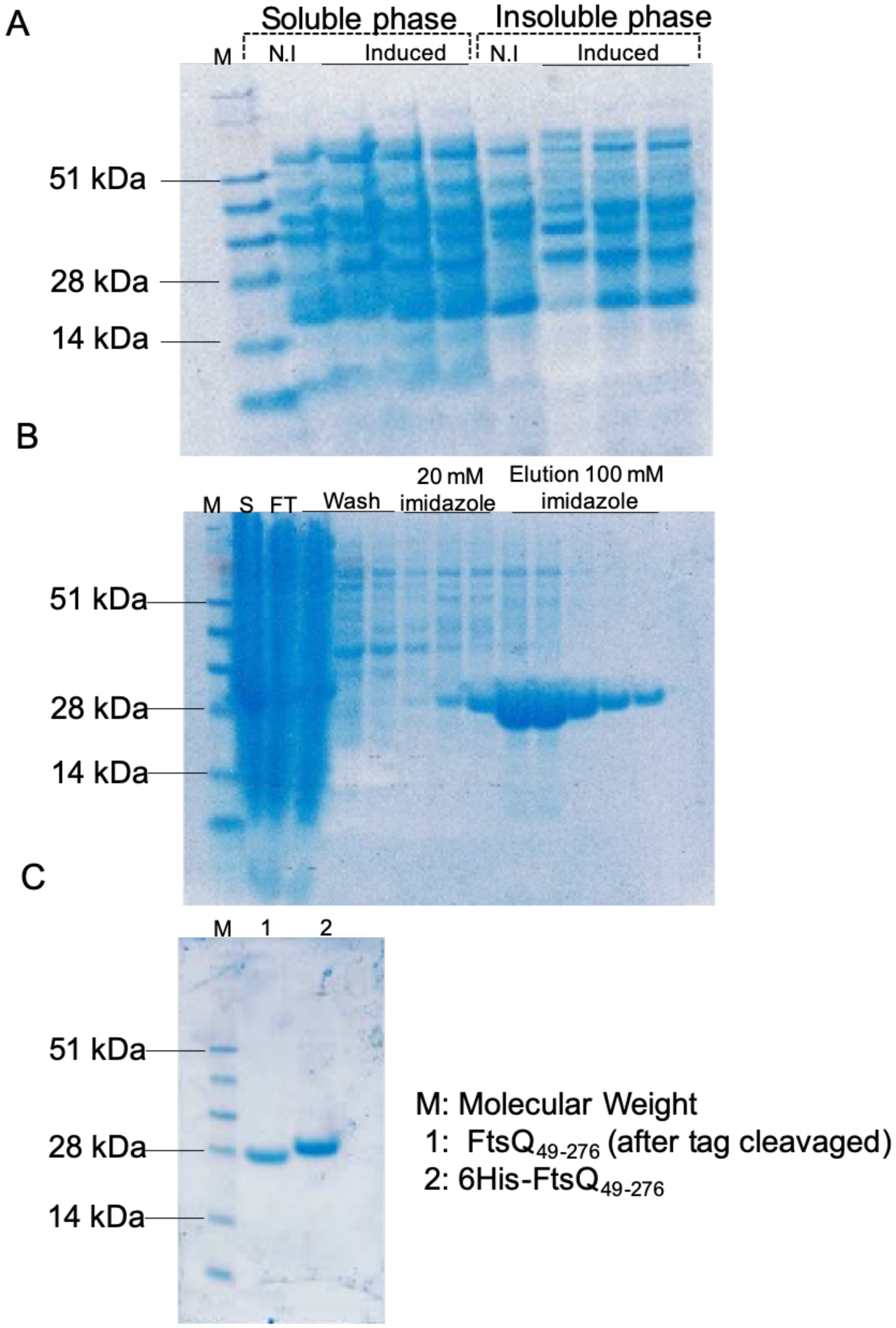


**Figure S1***: SDS-page analysis of 6His-FtsQ$_{49\text{-}276}$ production and purification. The proteins were separated by electrophoresis on a 4–12% SDS-polyacrylamide gel (Invitrogen). After migration, the gel was stained with PageBlue protein staining solution (Thermo Fisher Scientific)*. ***(A)*** *Analysis of total protein extract (soluble and insoluble fraction) after either early or late induction.* ***(B)*** *Analysis of protein purification fraction with Ni-NTA column.* ***(C)*** *Analysis of His-tag cleavage by the TEV protease*

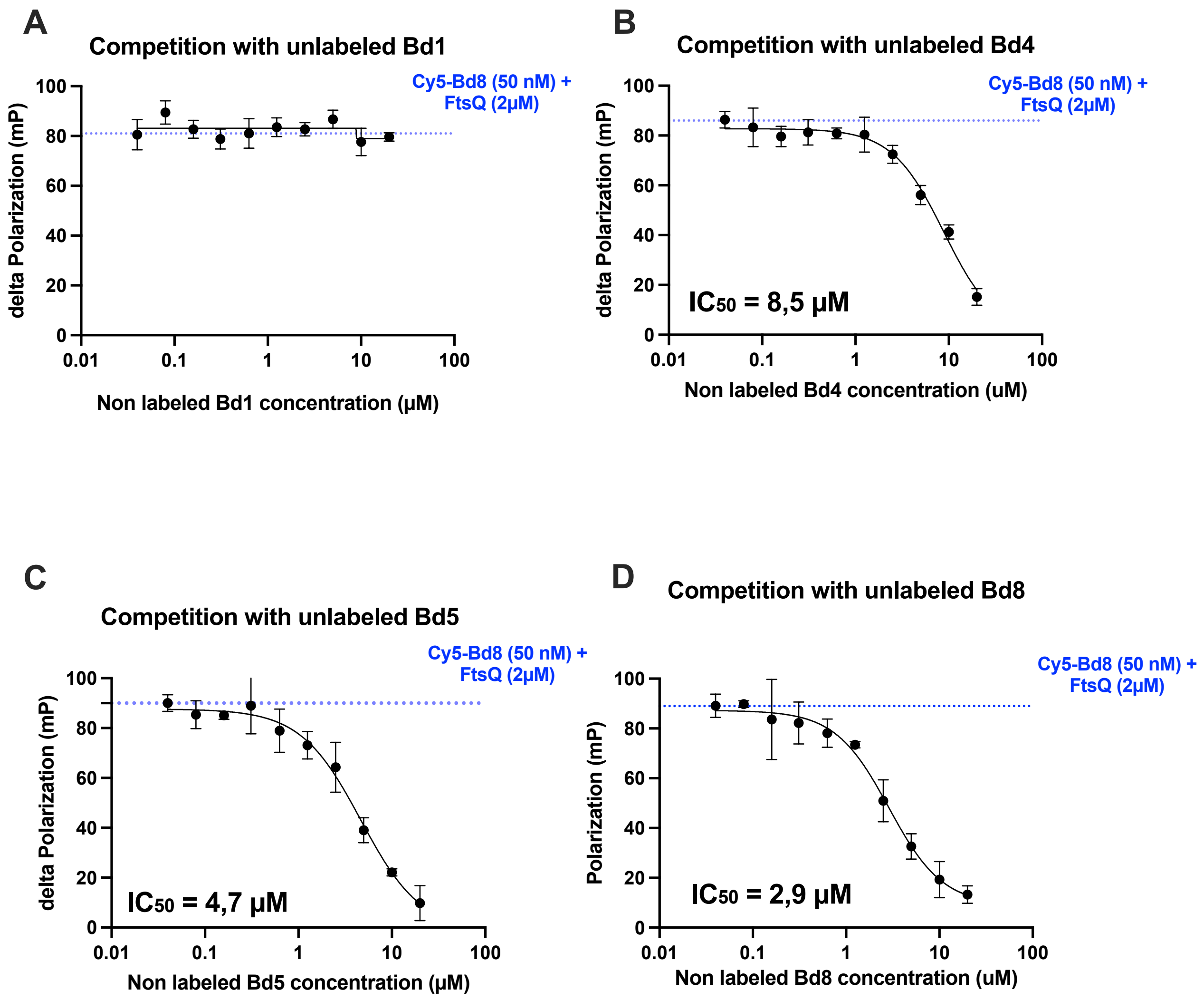


**Figure S2:** *Fluorescence polarization competition assay using Biotin tagged peptides Fluorescence polarization (FP) competition assays measuring displacement of Cy5–Bd8 by increasing concentrations of biotin tagged peptides (Bd1, Bd4, Bd5, Bd8). Assays were performed using 50 nM Cy5–Bd8 and 1.5 µM 6His–FtsQ$_{49–276}$. Data points and error bars represent the mean of three independent measurements and standard deviation. Data were fitted using a variable-slope dose–response inhibition model in Prism software.*

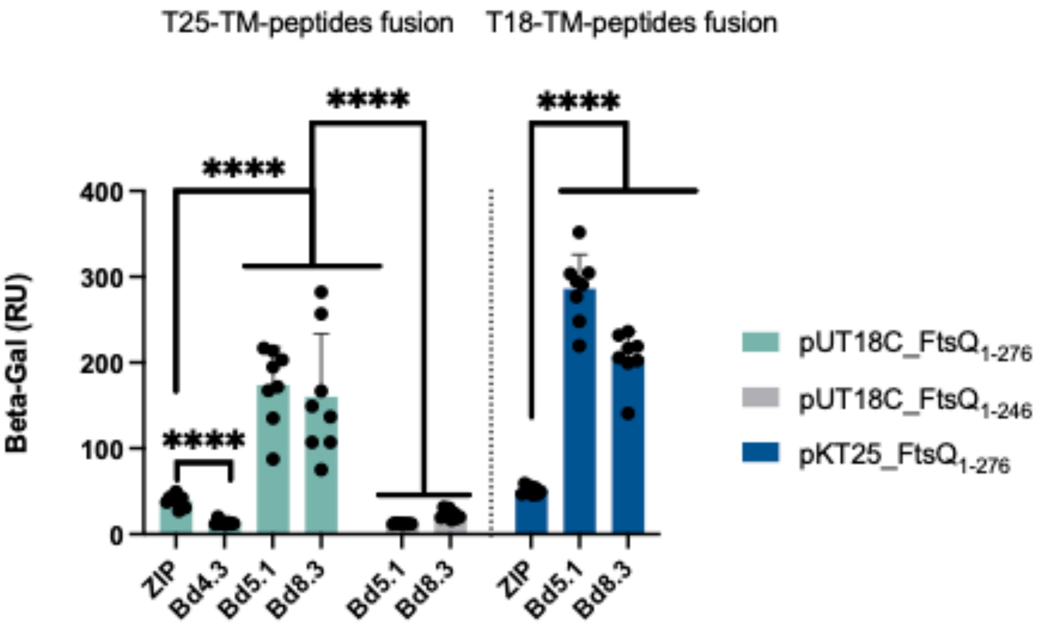


**Figure S3:** *Mapping of optimized peptides interaction with FtsQ proteins Functional complementation between the indicated proteins was assessed by measuring β-galactosidase activity in suspensions of chloroform-treated Escherichia coli cells harboring the corresponding plasmids, as described in Materials and Methods. Each bar represents the median of at least five independent cultures, and error bars indicate the interquartile range. A leucine zipper (ZIP) motif was used as a negative control. Periplasmic BACTH assay showing complementation between T25–transmembrane (TM) peptide fusions (pKTM25) and either full-length T18–FtsQ (residues 1–276; pUT18C-FtsQ1–276, light blue) or a C-terminally truncated variant (residues 1–246; pUT18C-FtsQ1–246, grey) in E. coli DHM1 cells. The interactions were also characterized using T18–transmembrane (TM) peptide fusions (pUTM18C) and full-length T25–FtsQ.*

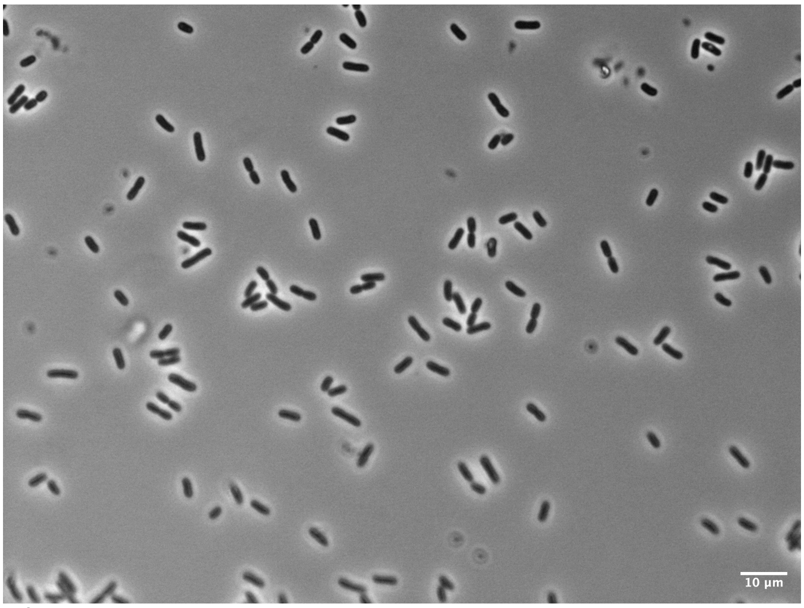


**Figure S4:** *Impact of peptide Bd4.3 overproduction on bacterial division:* *Phase-contrast microscopy images of Escherichia coli MG1655 cells overproducing T18–TM–Bd4.3 peptides (plasmid pUTM18C-Bd4.3, after 6 h 30 min of induction with 0.5 mM IPTG). Images were acquired at ×100 magnification. Scale bar 10µm*

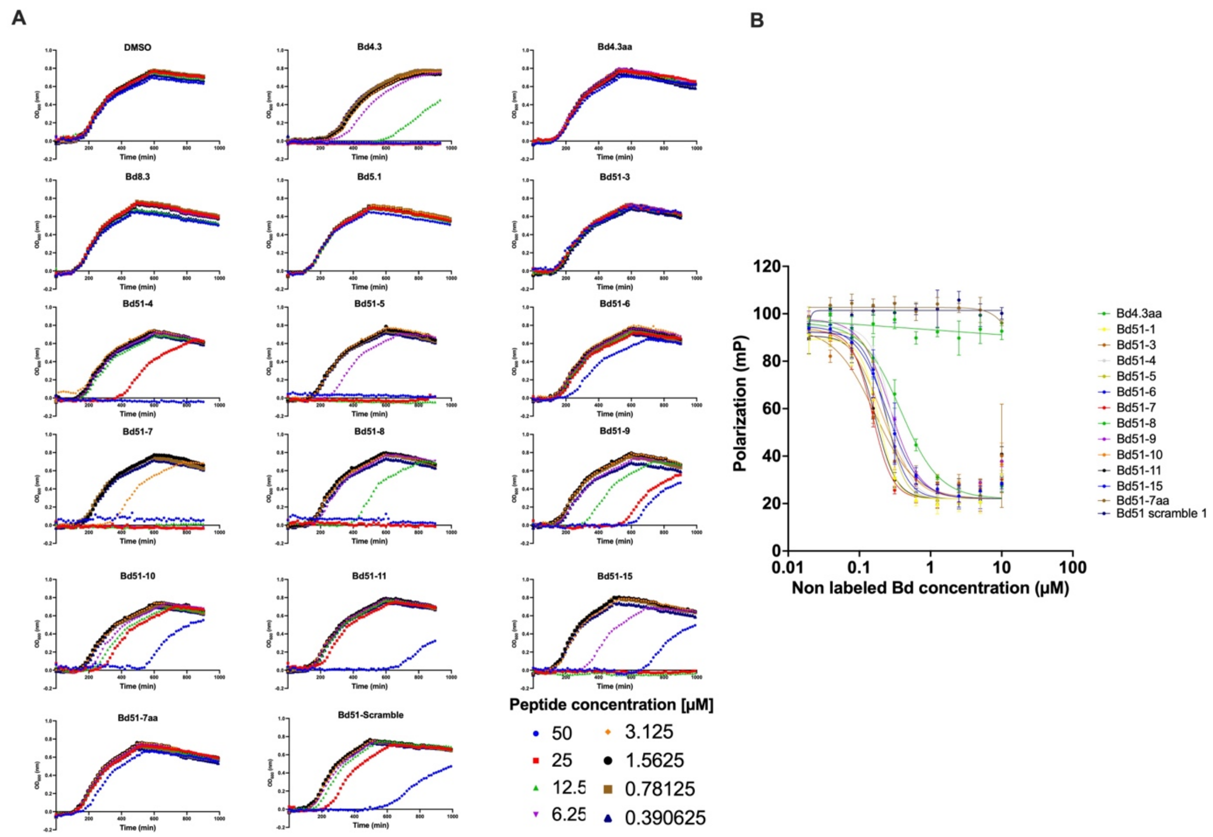


**Figure S5**: *Characterization of Bd51 derivatives peptides* ***(A)*** *Growth of E. coli lptD4213 in the presence of increasing concentrations of synthetic peptides or, as a control, equivalent concentrations of the solvent DMSO. Bacterial growth was monitored by measuring optical density at 600 nm ($OD_{600}$) in a microplate reader over 1000 min. Cultures were grown in LB medium at 37 °C with continuous agitation.* ***(B)*** *Fluorescence polarization (FP) competition assays measuring displacement of Cy5–Bd5 by increasing concentrations of unlabeled peptides. Assays were performed using 50 nM Cy5–Bd5 and 0.5 µM 6His–$FtsQ_{49–276}$. Data points represent the mean of two independent measurements, with error bars indicating the standard deviation. Data were fitted using a variable-slope dose–response inhibition model in Prism software.*

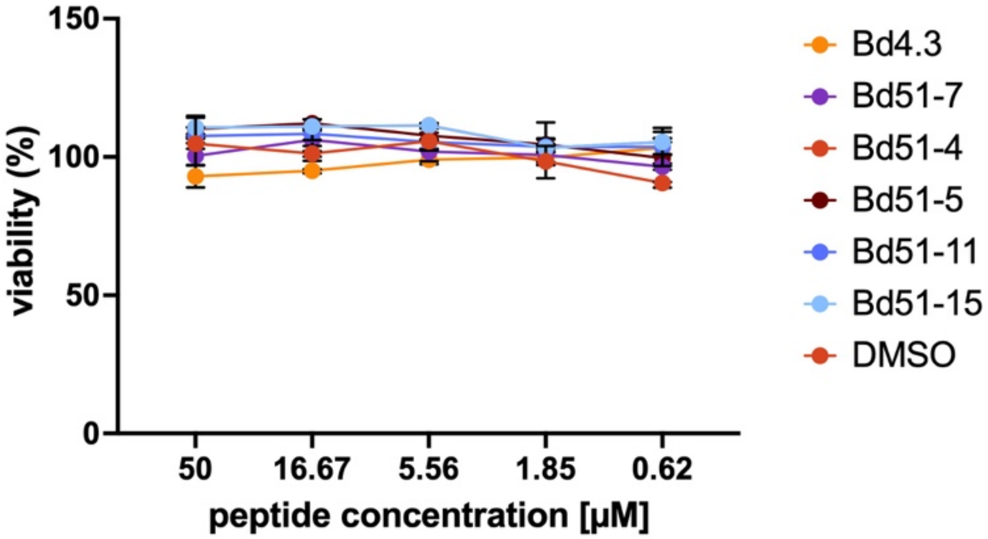


**Figure S6**: *Cytotoxicity measurement of Bd51 derivatives peptides Cytotoxicity assay on MRC5 human lung cells. Cell viability in presence of increasing concentration of peptide was assessed by Celltiter Glo viability assay and compared to untreated control cells (DMSO)l. Line represents best fit. Dots and error bars represent mean and standard deviation. n = 3 biologically independent samples.*

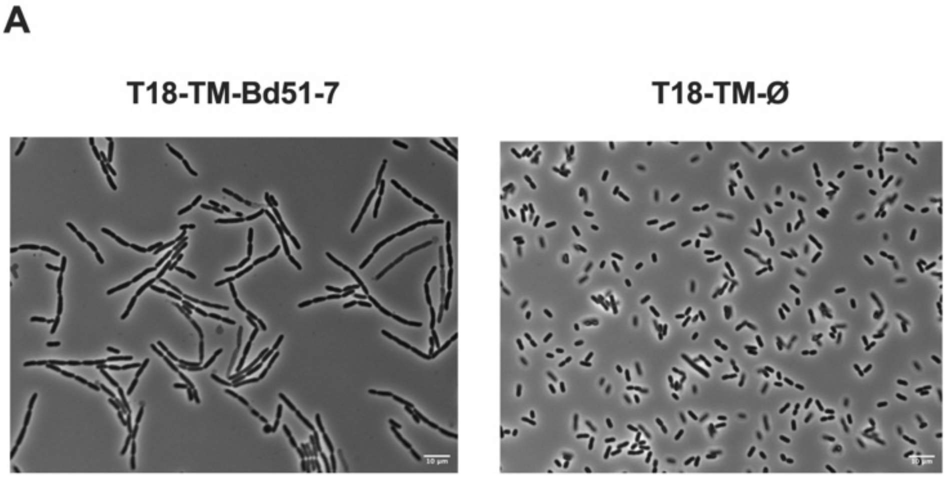


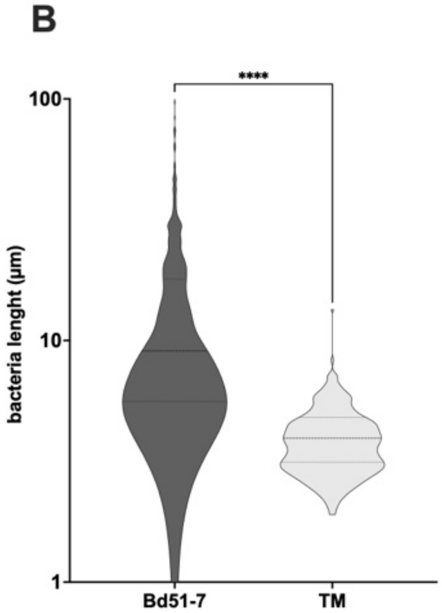


| | ø | Bd51-7 |
|---|---|---|
| Mean (µm) | 4.16 | 13.77 |
| Median (µm) | 3.94 | 9.07 |
| n (cells) | 224 | 544 |

**Figure S7:** *Impact of T18-TM-Bd51-7 peptide overproduction on bacterial division* ***(A)*** *Phase-contrast microscopy images of Escherichia coli MG1655 cells overproducing T18–TM–Bd51-7 peptides (plasmid pUTM18C-Bd51-7,after 6 h 30 min of induction with 0.5 mM IPTG). Images were acquired at ×100 magnification. Scale bar 10µm****(B)*** *Quantification of bacterial cell length following 6 h 30 min of peptide overexpression, measured using ObjectJ software. Median and interquartile range are showed. One way Brown-Forsythe and Welch Anova test with Games-Howell Post hoc, ****P < 0.0001*

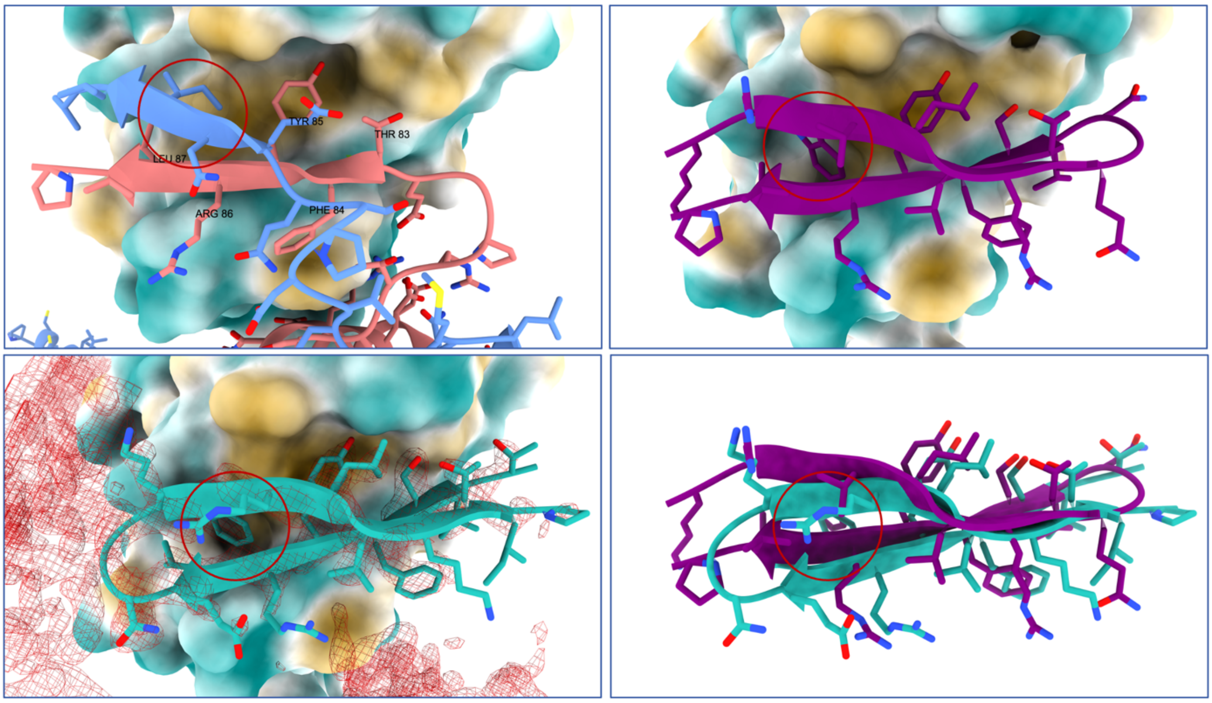


***Figure S8:*** *Structural comparison between native and designed peptide interactions at the FtsQ interface.* *(A) Native FtsQBL complex (PDB: 8HHH) showing β-strand augmentation of FtsQ by FtsB/FtsL at the divisome interface. Hotspot residues on FtsQ used as positional constraints during RFdiffusion-guided design are indicated. (B) Crystal structure of the designed peptide Bd4.3 bound to FtsQ, reproducing the native β-strand interaction geometry while introducing a conserved tryptophan residue that inserts into the same hydrophobic pocket. (C) Experimental electron density supporting the positioning of Bd4.3 at the interface and confirming the orientation of the tryptophan side chain within the pocket. (D) AlphaFold model of the Bd5.1-7 peptide (magenta) superimposed onto the Bd4.3 crystal structure (cyan). Although Bd5.1-7 contains a loop extension positioned on the opposite side of the interface, the model predicts conserved interactions along the FtsQ β-sheet and preservation of pocket engagement by the conserved tryptophan residue (red circle), corresponding to the position occupied by an isoleucine from FtsL in the native complex.*

**Table S1: Peptides sequences**

| Name | Sequence | Method of Design | BACTH dosage (R.U)[1] |
|---|---|---|---|
| Bd 1 | INEQSADVPGETFYRLVP | Based on native sequence FtsB/FtsL | 7 |
| Bd 2 | INEQCADVPGECFYRLVP | Based on native sequence FtsB/FtsL | N.A |
| Bd 3 | INEQSADCPGCTFYRLVP | Based on native sequence FtsB/FtsL | 6 |
| Bd 4 | TETFYRLEDGEWSLVRTVE | RF diffusion | 177 |
| Bd4.1 | TVSFYRLVDGKWVLVRTVP | RF diffusion based on Binder 4 sequence | 177 |
| Bd4.2 | TVSFYRLVDGKWVHVRTVP | RF diffusion based on Binder 4 sequence | 198 |
| Bd4.3 | TVSFYRLENGKWRLVKTIP | RF diffusion based on Binder 4 sequence | 238 |
| Bd4.4 | TVSFYELRDGEWRLVRTVP | RF diffusion based on Binder 4 sequence | 375 |
| Bd4.5 | TVSFYELVDGEWRLVRTVP | RF diffusion based on Binder 4 sequence | 149 |
| Bd4.6 | TVTFYRLVDGEWVHVRTVP | RF diffusion based on Binder 4 sequence | 192 |
| Bd4.7 | TVSFYRLVDGEWVHVRTVP | RF diffusion based on Binder 4 sequence | 216 |
| Bd4.8 | TVSFYRLEDGEWVLVSTVP | RF diffusion based on Binder 4 sequence | 195 |
| Bd4.9 | TVSFYELVDGEWRLVSTVP | RF diffusion based on Binder 4 sequence | 145 |
| Bd4.10 | TVSFYELRDGEWRLVSTVP | RF diffusion based on Binder 4 sequence | 155 |
| Bd 5 | TRWERVRTEGDRTFYRLVE | RF diffusion | 59 |
| Bd5.1 | ERWVLVRTEGDVSFYELVP | RF diffusion based on Binder 5 sequence | 164 |
| Bd5.2 | SRWVKVRTEGDVTFYELVP | RF diffusion based on Binder 5 sequence | 160 |
| Bd5.3 | TRWVHVRTEGDVTFYEQVP | RF diffusion based on Binder 5 sequence | 81 |
| Bd5.4 | TRWVLVRTEGDVSYYEEVP | RF diffusion based on Binder 5 sequence | 198 |
| Bd5.5 | TRWVLVSTEGDVSFYEEVP | RF diffusion based on Binder 5 sequence | 260 |
| Bd5.6 | TEWRLVSTEGDVSFYEEVP | RF diffusion based on Binder 5 sequence | 123 |
| Bd5.7 | SVWVKVRTEGDVTFYEEVP | RF diffusion based on Binder 5 sequence | 107 |
| Bd5.8 | SRWVHVRTEGDVTFYREVP | RF diffusion based on Binder 5 sequence | 297 |
| Bd5.9 | TVWVKVSTEGDVTFYEEVP | RF diffusion based on Binder 5 sequence | 257 |
| Bd5.10 | TRWVLVSTEGDESFYEEVP | RF diffusion based on Binder 5 sequence | 165 |
| Bd 6 | TRWERVRCEGDRCFYRLVE | RF diffusion | 27 |
| Bd 7 | STRSWEAVRREGDKTFYRLVEN | RF diffusion | 51 |
| Bd 8 | AARWEAVRTEGDRTFYRLLPPE | RF diffusion | 358 |
| $Bd8_{F15A}$ | AARWEAVRTEGDRTAYRLLPPE | Alanine point mutation in Bd8 F15A | 14 |
| $Bd8_{Y16A}$ | AARWEAVRTEGDRTFARLLPPE | Alanine point mutation in Bd8 Y16A | 14 |
| $Bd8_{AA}$ | AARWEAVRTEGDRTAARLLPPE | Alanine point mutation in Bd8 FY15/16AA | 14 |
| Bd8.1 | AAQWELVSTEGDVSLYRELPAP | RF diffusion based on Binder 8 sequence | 110 |
| Bd8.2 | KPQWVLVSTEGDVSFYVEQPAP | RF diffusion based on Binder 8 sequence | 274 |
| Bd8.3 | KAKWVLVRTEGDVSYYEEQPAP | RF diffusion based on Binder 8 sequence | 206 |
| Bd8.4 | KPQWVLVSTEGDVSHYVEQPAP | RF diffusion based on Binder 8 sequence | 262 |
| Bd8.5 | KPQWRLVRTEGDVSFYEEQPAP | RF diffusion based on Binder 8 sequence | 193 |

| | | | |
|---|---|---|---|
| Bd8.6 | AAQWELVETRGDESLYRELPAP | RF diffusion based on Binder 8 sequence | 152 |
| Bd8.7 | KAQWRLVRTEGDVSLYEELPAP | RF diffusion based on Binder 8 sequence | 96 |
| Bd8.8 | KPQWELVRTEGDVSLYRELPAP | RF diffusion based on Binder 8 sequence | 97 |
| Bd8.9 | KPQWELVRTEGDVSLYRELPAP | RF diffusion based on Binder 8 sequence | 122 |
| Bd8.10 | KPQWRLVSTEGDVSLYEELPAP | RF diffusion based on Binder 8 sequence | 11 |
| Bd51-1 | SRWVLVRTEGDVSFYRLVP | Rational design based on Bd5.1 sequence | N.A |
| Bd51-3 | SRWVLVRTEPDVSFYRLVP | Rational design based on Bd5.1 sequence | N.A |
| Bd51-4 | SRWVLVRTEGNVSFYRLVP | Rational design based on Bd5.1 sequence | N.A |
| Bd51-5 | SRWVLVRTQGNVSFYRLVP | Rational design based on Bd5.1 sequence | N.A |
| Bd51-6 | KRWVLVRTEGDVSFYRLVP | Rational design based on Bd5.1 sequence | N.A |
| Bd51-7 | KRWVLVRTQGNVSFYRLVP | Rational design based on Bd5.1 sequence | N.A |
| Bd51-8 | KRWVLVRTQPNVSFYRLVP | Rational design based on Bd5.1 sequence | N.A |
| Bd51-9 | RWVLVRTEGNVSFYRL | Rational design based on Bd5.1 sequence | N.A |
| Bd51-10 | WVLVRTQGNVSFYRL | Rational design based on Bd5.1 sequence | N.A |
| Bd51-11 | QRWVLVRTEGDVSFYRLVP | Rational design based on Bd5.1 sequence | N.A |
| Bd51-15 | RWVLVRTQGNVSFYRL | Rational design based on Bd5.1 sequence | N.A |
| Bd51-$7_{AA}$ | KRWVLVRTQGNVSAARLVP | Alanine point mutations F14A & Y15A | N.A |
| Bd51-Scramble | TFRSVLNVQRYGVLPKVWR | Shuffled sequence of Bd5.1 | N.A |
| BdP.1[2] | RETLYQLVDGEVVASPL | RF diffusion | 64 |
| BdP.2[2] | METLYQLQPDGTVVASPL | RF diffusion | 144 |
| BdP.3[2] | KETLYQLNPDGTVTASE | RF diffusion | 18 |
| BdP.4[2] | KETLYQYNEETGEVTASPL | RF diffusion | 111 |
| BdP.5[2] | METLYQLEGDEVVASPL | RF diffusion | 14 |
| BdP.6[2] | ITTYLSLDEPEEELLAKVAAALA AVAAA | RF diffusion | 14 |
| BdP.7[2] | KTTYYDELGLE | RF diffusion | 14 |
| BdP.8[2] | MEYITEERAEELE | RF diffusion | 14 |
| BdP.9[2] | SVIDEAVERIKKEIAERAAK | RF diffusion | 14 |
| BdP.10[2] | AKYVEALITEENAEEAVAAAKAA VK | RF diffusion | 14 |

[1]BACTH dosage (R.U) correspond to the median of 8 independent ß-galactosidase dosage made in E. coli DHT1 cells using either T25-Bd and T18-$FtsQ_{49-276}$ partners or T25-BdP and T18-$FtsQ_{P.A65-285}$

[2] Peptides designed to target FtsQ homologous protein from *Pseudomonas aeruginosa*

-

### Table S2: Sequence optimization of RFdiffusion-derived peptides Binder4, Binder5, and Binder8 targeting the FtsQ periplasmic groove

| | FtsL (Cter->Ner) | | | | | | | | | | FtsB (Cter->Ner) | | | | | | | | | |
|---|---|---|---|---|---|---|---|---|---|---|---|---|---|---|---|---|---|---|---|---|
| Native sequence | V | V | I | N | E | Q | S | P | D | V | P | G | E | T | F | Y | R | L | V | P |
| Residue ID | 119 | 118 | 117 | 116 | 115 | 114 | 113 | 112 | 111 | 110 | 89 | 88 | 87 | 86 | 85 | 84 | 83 | 82 | 81 | 80 |

| | | Hotspots |
|---|---|---|
| FtsB | Nter -> Cter | P G E T F Y R L V P |
| FtsL | Nter -> Cter | V D P S Q E N I V V |

| Design | | | | | N-terminal | | | | | | | Loop | | | | C-terminal | | | | | | | | | PAE interaction | Domain layout |
|---|---|---|---|---|---|---|---|---|---|---|---|---|---|---|---|---|---|---|---|---|---|---|---|---|---|---|
| | | 1 | 2 | 3 | 4 | 5 | 6 | 7 | 8 | 9 | 10 | 11 | 12 | 13 | 14 | 15 | 16 | 17 | 18 | 19 | 20 | 21 | 22 | 23 | | |
| RF-diffusion | Binder4 | | | | T | E | T | F | Y | R | L | E | D | G | E | W | S | L | V | R | T | V | E | | 5.279 | FtsB-mimic N-ter -> T E T F Y R L E D; FtsL-mimic C-ter <- E V T R V L S W E G |
| | | | | | | ↓ | ↓ | | | | | | ↓ | | ↓ | | ↓ | | | ↓ | | ↓ | ↓ | | | |
| RF-diffusion | Binder4.3 | | | | T | V | S | F | Y | R | L | E | N | G | K | W | R | L | V | K | T | I | P | | 4.574 | FtsB-mimic N-ter -> T V S F Y R L E N; FtsL-mimic C-ter <- P I T K V L R W K G |
| | | | | | | | | | | ↓ | | ↓ | | | | | ↓ | | | | | | ↓ | | | |
| RF-diffusion | Binder4.4 | | | | T | V | S | F | Y | E | L | R | D | G | E | W | R | L | V | R | T | V | P | | 4.582 | FtsB-mimic N-ter -> T V S F Y E L R D; FtsL-mimic C-ter <- P V T R V L R W E G |
| RF-diffusion | Binder5 | | | T | R | W | E | R | V | R | T | E | G | D | R | T | F | Y | R | L | V | E | | | 5.346 | FtsB-mimic -> C-ter D R T F Y R L V E; FtsL-mimic <- N-ter G E T R V R E W R T |
| | | | | ↓ | | | ↓ | ↓ | | | | | | | ↓ | ↓ | | | ↓ | | | ↓ | | | | |
| RF-diffusion | Binder5.1 | | | E | R | W | V | L | V | R | T | E | G | D | V | S | F | Y | E | L | V | P | | | 4.852 | FtsB-mimic -> C-ter D V S F Y E L V P; FtsL-mimic <- N-ter G E T R V L V W R E |
| | | | | ↓ | | | ↓ | ↓ | | | | | | | ↓ | | | | ↓ | | | ↓ | | | | |
| RF-diffusion | Binder5.2 | | | S | R | W | V | K | V | R | T | E | G | D | V | T | F | Y | E | L | V | P | | | 5.002 | FtsB-mimic -> C-ter D V T F Y E L V P; FtsL-mimic <- N-ter G E T R V K V W R S |
| | | | | | | | ↓ | ↓ | | | | | | | ↓ | ↓ | ↓ | | ↓ | ↓ | | ↓ | | | | |
| RF-diffusion | Binder5.4 | | | T | R | W | V | L | V | R | T | E | G | D | V | S | Y | Y | E | E | V | P | | | 5.141 | FtsB-mimic -> C-ter D V S Y Y E E V P; FtsL-mimic <- N-ter G E T R V L V W R T |
| | | | | | | | ↓ | ↓ | | ↓ | | | | | ↓ | ↓ | | | ↓ | ↓ | | ↓ | | | | |
| RF-diffusion | Binder5.5 | | | T | R | W | V | L | V | S | T | E | G | D | V | S | F | Y | E | E | V | P | | | 5.168 | FtsB-mimic -> C-ter D V S F Y E E V P; FtsL-mimic <- N-ter G E T S V L V W R T |
| | | | | | | | ↓ | ↓ | | ↓ | | | | | ↓ | ↓ | | | ↓ | ↓ | | ↓ | | | | |
| RF-diffusion | Binder5.10 | | | T | R | W | V | L | V | S | T | E | G | D | E | S | F | Y | E | E | V | P | | | 5.312 | FtsB-mimic -> C-ter D V S F Y E E V P; FtsL-mimic <- N-ter G E T S V L V W R T |
| RF-diffusion | Binder8 | | A | A | R | W | E | A | V | R | T | E | G | D | R | T | F | Y | R | L | L | P | P | E | 6.503 | FtsB-mimic -> C-ter D R T F Y R L L P P E; FtsL-mimic <- N-ter G E T R V A E W R A A |
| | | | ↓ | ↓ | ↓ | | ↓ | ↓ | | ↓ | | | | | ↓ | ↓ | | | ↓ | ↓ | ↓ | | ↓ | ↓ | | |
| RF-diffusion | Binder8.2 | | K | P | Q | W | V | L | V | S | T | E | G | D | V | S | F | Y | V | E | Q | P | A | P | 6.308 | FtsB-mimic -> C-ter D V S F Y V E Q P A P; FtsL-mimic <- N-ter G E T S V L V W Q P K |
| | | | ↓ | | ↓ | | ↓ | ↓ | | | | | | | ↓ | ↓ | ↓ | | ↓ | ↓ | ↓ | | ↓ | ↓ | | |
| RF-diffusion | Binder8.3 | | K | A | K | W | V | L | V | R | T | E | G | D | V | S | Y | Y | E | E | Q | P | A | P | 6.384 | FtsB-mimic -> C-ter D V S Y Y E E Q P A P; FtsL-mimic <- N-ter G E T R V L V W K A K |
| | | | ↓ | ↓ | ↓ | | ↓ | ↓ | | | | | | | ↓ | ↓ | | | ↓ | ↓ | ↓ | | ↓ | ↓ | | |
| RF-diffusion | Binder8.5 | | K | P | Q | W | R | L | V | R | T | E | G | D | V | S | F | Y | E | E | Q | P | A | P | 6.473 | FtsB-mimic -> C-ter D V S F Y E E Q P A P; FtsL-mimic <- N-ter G E T R V L R W Q P K |

| Legend | |
|---|---|
| | N-ter |
| | Loop |
| | C-ter |

*For each peptide, the original design and optimized variants are aligned along the FtsB/FtsL β-augmentation register. The vertical arrows indicate point mutations introduced between successive design rounds. These substitutions were selected to improve the predicted interface geometry and stability as evaluated by the AlphaFold2 PAE_interaction score, shown on the right of each sequence block. Lower PAE_interaction values correspond to higher confidence in the modeled peptide–FtsQ interaction. While the TFYRL β-strand recognition motif is conserved across binder 4, 5, and 8, only Y85 is preserved in all optimized peptides underscoring its essential role in maintaining the native FtsB/L-like β-augmentation within the FtsQ groove.*

### Table S3 Bd4.3 - Chemical shifts*

| Residue | Atom | H/N | HA/CA | HB/CB | HG/CG | HD/CD | HE/CE | Other |
|---|---|---|---|---|---|---|---|---|
| 1Thr | H | 8.15 | 4.42 | 4.1 | 1.16 | | | Acetyl: 2.08 |
| | X | 120.4 | 62.2 | 70.3 | 21.8 | | | Acetyl: 24.5 |
| 2Val | H | 8.39 | 4.34 | 1.98 | 0.77,0.89 | | | |
| | X | 123.7 | 63.1 | 33.6 | 20.63,21.53 | | | |
| 3Ser | H | 8.27 | 4.91 | 3.65,3.65 | | | | |
| | X | 119.9 | | 65.2 | | | | |
| 4Phe | H | 8.50 | 5.00 | 2.90,3.04 | | 7.04 | 7.20 | Hz 7.21 |
| | X | 121.2 | | 40.8 | | 131.9 | 131.2 | Cz 129.9 |
| 5Tyr | H | 7.21 | 5.33 | 2.78,2.94 | | 6.87 | 6.71 | |
| | X | | 57.4 | 42.0 | | 133.0 | 118.3 | |
| 6Arg | H | 9.19 | 4.57 | 1.70,1.72 | 1.47,1.47 | 3.11,3.11 | 7.37 | |
| | X | | | | 27.0 | | | |
| 7Leu | H | 8.54 | 4.11 | 1.33 | 0.82 | -0.10,0.40 | | |
| | X | | 54.6 | | | 24.7 | | |
| 8Glu | H | 8.84 | 4.46 | 1.84,1.93 | 2.12,2.18 | | | |
| | X | 126.6 | 55.7 | 32.1 | 36.2 | | | |
| 9Asn | H | 7.62 | 4.36 | 2.74,3.01 | | 6.92,7.60 | | |
| | X | | 54.1 | 37.7 | | 112.8 | | |
| 10Gly | H | 8.36 | 3.46,4.06 | | | | | |
| | X | 103.3 | 45.7 | | | | | |
| 11Lys | H | 7.63 | 4.56 | 1.74,1.81 | 1.39,1.39 | 1.70,1.70 | 3.02,3.02 | |
| | X | 120.4 | 54.5 | 35.0 | 24.8 | 29.0 | 42.3 | |
| 12Trp | H | 8.63 | 4.86 | 3.01,3.05 | | 7.20 | 7.21 | He1: 9.99 Hh: 7.34 Hz: 7.04 |
| | X | 123.8 | | 29.9 | | 127.4 | 119.7 | |
| 13Arg | H | 9.3 | 4.77 | 1.79,1.86 | 1.61,1.65 | 3.18,3.18 | 7.27 | |
| | X | | | 32.9 | 27.0 | 43.5 | | |
| 14Leu | H | 8.64 | 4.04 | 1.35,1.61 | 1.23 | 0.56,0.72 | | |
| | X | 126.7 | 55.4 | 42.3 | 27.0 | 23.52,25.25 | | |
| 15Val | H | 8.86 | 4.2 | 1.83 | 0.79,0.85 | | | |
| | X | | 62.0 | 33.5 | 20.60,21.44 | | | |
| 16Lys | H | 8.11 | 4.58 | 1.73,1.79 | 1.30,1.37 | 1.59,1.59 | 2.84 | |
| | X | | 56.0 | 34.2 | 24.8 | 29.3 | 42.0 | |
| 17Thr | H | 8.24 | 4.61 | 4.03 | 1.13 | | | |
| | X | | | 70.5 | 21.6 | | | |
| 18Ile | H | | | 1.85 | 0.87,1.04,1.43 | 0.82 | | |
| | X | | | 39.0 | 17.2 | 13.1 | | |
| 19Pro | H | | 4.36 | 1.90,2.26 | 2.01,2.02 | 3.56,3.78 | | NH2: 6.93,6.70 |
| | X | | 61.9 | 32.3 | 27.4 | 49.8 | | NH2: 107.6 |

**: $^{1}$H, $^{13}$C and $^{15}$N chemical shifts of Bd4.3 obtained at 25 °C. The N-terminal-acylated and C-terminal-amidated peptide (517 µM) was dissolved in 20 mM sodium phosphate pH 6.2 and 3% $D_2O$. Residues Y5, R6, L7, I18 and P19 show broad lines due to conformational exchange in the µs-ms time range. Residue P19 is in a trans-conformation as determined by the CB chemical shift. There is a minor population of cis conformer that produces additional spin systems for Thr1 and Ile18 that are close in space. Chemical shifts are referenced to DSS (sodium 2,2-dimethyl-2-silapentane-5-sulfonate).*

### Table S4 Bd5.1-7 – Chemical shifts*

| Residue | Atom | H/N | HA/CA | HB/CB | HG/CG | HD/CD | HE/CE | Other |
|---|---|---|---|---|---|---|---|---|
| 1Lys | H | 8.18 | 4.32 | 1.40,1.55 | 1.28,1.28 | 1.59,1.59 | 2.93 | Acetyl: 2.03 |
| | X | 126.5 | 56.4 | 33.7 | 25.2 | 29.4 | 42.3 | Acetyl: 24.5 |
| 2Arg | H | 8.02 | 4.26 | 1.69,1.78 | 1.54,1.54 | 3.19 | | |
| | X | | 55.8 | 31.3 | 27.2 | 43.6 | | |
| 3Trp | H | 8.36 | 5.09 | 3.04,3.06 | | 7.13 | 7.35 | He1: 10.00 Hh2: 7.17 Hz2: 7.42Hz3: 7.05 |
| | X | 122.3 | 56.6 | 30.8 | | 127.3 | 120.4 | Ne1: 129.3 Ch2: 124.6 Cz2: 114.8 Cz3: 122.0 |
| 4Val | H | | 4.48 | 2.02 | 0.89,0.92 | | | |
| | X | | | 34.7 | 20.3,21.0 | | | |
| 5Leu | H | 8.47 | 4.09 | 1.25,1.38 | 1.63 | 0.57,0.74 | | |
| | X | 127.2 | 55.4 | 42.4 | 27.1 | 23.7,25.1 | | |
| 6Val | H | 8.15 | 4.45 | 2.11 | 0.93,0.97 | | | |
| | X | 122.9 | 59.6 | 32.9 | 20.3,21.1 | | | |
| 7Arg | H | 8.09 | 4.65 | 1.74,1.84 | 1.49,1.49 | 2.95,3.09 | | |
| | X | | 55.6 | 31.8 | 27.1 | 43.4 | | |
| 8Thr | H | 8.45 | 4.71 | 4.19 | 1.19 | | | Hg1: 5.61 |
| | X | 116.1 | | 70.6 | 21.7 | | | |
| 9Gln | H | 8.73 | 4.41 | 2.03,2.15 | 2.37,2.38 | | 6.88,7.55 | |
| | X | 123.7 | 56.1 | 29.8 | 33.9 | | 112.5 | |
| 10Gly | H | 8.73 | 3.88,3.98 | | | | | |
| | X | 112.3 | 46.2 | | | | | |
| 11Asn | H | | 4.74 | 2.77,2.83 | | 6.91,7.57 | | |
| | X | | 53.1 | 39.1 | | 112.7 | | |
| 12Val | H | 7.87 | 4.20 | 2.09 | 0.78,0.93 | | | |
| | X | 120.4 | 62.4 | 33.0 | 21.5,21.0 | | | |
| 13Ser | H | 8.38 | 4.85 | 3.59,3.67 | | | | |
| | X | 121.2 | | 64.9 | | | | |
| 14Phe | H | 8.39 | 4.76 | 2.94,3.08 | | 7.10 | 7.25 | Hz: 7.28,7.26 |
| | X | 120.7 | | 40.6 | | 132.0 | 131.3 | Cz: 130.0 |
| 15Tyr | H | 8.38 | 5.14 | 2.77,2.93 | | 6.92 | 6.73 | |
| | X | 120.8 | | 40.6 | | 133.0 | 118.4 | |
| 16Arg | H | 8.36 | 4.48 | 1.68,1.74 | 1.50 | 3.14,3.14 | | |
| | X | | 55.4 | 32.0 | 27.1 | 43.5 | | |
| 17Leu | H | 8.39 | 4.30 | 0.97,1.37 | 1.05 | 0.36,0.51 | | |
| | X | 126.0 | 54.74 | 42.4 | 26.7 | 23.7,24.4 | | |
| 18Val | H | 8.61 | 4.55 | 2.12 | 0.89,0.94 | | | |
| | X | 124.4 | 59.35 | 32.9 | 20.3,21.1 | | | |
| 19Pro | H | | 4.33 | 1.94,2.29 | 2.07,2.07 | 3.70,3.83 | | NH2:7.61, 6.99 |
| | X | | 63.2 | 32.4 | 27.5 | 51.0 | | NH2: 107.5 |

*: *$^{1}$H, $^{13}$C and $^{15}$N chemical shifts of Bd5.1-7 obtained at 25 °C. The N-terminal-acylated and C-terminal-amidated peptide (667 µM) was dissolved in 10 mM sodium phosphate, 20 mM deuterated TRIS-d18 pH 6.3, 150 mM NaCl and 3% $D_2O$. Residue P19 is in a trans-conformation as determined by the V18HA-P19HD* nOe with the preceding residue. There is a second population that shows different chemical shifts for the aromatic rings of F14 and Y15 and random coil chemical shifts for those spin systems that could be partially assigned. Chemical shifts are referenced to DSS (sodium 2,2-dimethyl-2-silapentane-5-sulfonate).*

**Table S5: Plasmids**

| Name | Vector features | Characteristic | Reference |
|---|---|---|---|
| pKT25 | Kanamycin p15A Ori | Empty vector, encodes the T25 fragment of CyaA | Karimova et al., 2001 |
| pUT18C | Ampicillin colE1 Ori | Empty vector, encodes the T18 fragment of CyaA | Karimova et al., 2001 |
| pUTM18C | Kanamycin p15A Ori | Empty vector, encodes the T25 fragment of CyaA fused to a TM domain | Ouellette et al. 2013 |
| pKTM25 | Ampicillin colE1 Ori | Empty vector, encodes the T18 fragment of CyaA fused to a TM domain | Ouellette et al. 2013 |
| pKT25-ZIP | Kanamycin p15A Ori | Encodes the T25 fragment of CyaA fused to a leucine zipper | Karimova et al., 2001 |
| pUT18C-ZIP | Ampicillin colE1 Ori | Encodes the T18 fragment of CyaA fused to a leucine zipper | Karimova et al., 2001 |
| pKTM25-BdX$_{reftable}$ | Kanamycin p15A Ori | Encodes the T25 fragment of CyaA fused to OppB TM domain and to a design peptide (cf peptide table) under the control of a lac promoter | This work |
| pKT25- BdX$_{reftable}$ | p15A Ori | Encodes the T25 fragment of CyaA fused to a design peptide (cf peptide table) under the control of a lac promoter | This work |
| pUT18C-FtsQ$_{1-276}$ | Ampicillin colE1 Ori | Encodes the T18 fragment of CyaA fused to the FtsQ protein under the control of a lac promoter | Karimova et al. 2005 |
| pUT18C-FtsQ$_{49-276}$ | Ampicillin colE1 Ori | Encodes the T18 fragment of CyaA fused to the periplasmic domain of FtsQ protein under the control of a lac promoter | This work |
| pUT18C-FtsQ$_{1-246}$ | Ampicillin colE1 Ori | Encodes the T18 fragment of CyaA fused to the FtsQ protein without C-terminal extremity under the control of a lac promoter | Karimova et al. 2005 |
| pUT18C-FtsQ$_{49-246}$ | Ampicillin colE1 Ori | Encodes the T18 fragment of CyaA fused to the periplasmic domain of FtsQ protein without C-terminal extremity under the control of a lac promoter | This work |

| | | | |
|---|---|---|---|
| pUT18C-FtsQ$_{P.a65-287}$ | Ampicillin colE1 Ori | Encodes the T18 fragment of CyaA fused to the periplasmic domain of FtsQ protein from *P.aeruginosa* under the control of a lac promoter | This work |
| pKN-FtsQ | Kanamycin p15A Ori | Encodes the FtsQ protein | This work |
| pETM-FtsQ$_{49}$ | Kanamycin colE1 Ori | Express the 6His-FtsQ$_{49}$ protein under the control of a T7 promoter, a TEV cleavage site is present between the His tag and FtsQ$_{49}$ | This work |
| pKTM25-Bd8-T18FtsQ | Kanamycin p15A Ori | Encodes both the T25 fragment of CyaA fused to OppBTM-Bd8 peptide and the T18 fragment fused to FtsQ under the control of a lac promoter | This work |
| pUC19-FtsB | Ampicillin colE1 Ori | Encodes FtsB fused to T18 fragment of CyaA under the control of a lac promoter | Karimova et al., 2005 |
| pUC19-FtsL | Ampicillin colE1 Ori | Encodes FtsL fused to T18 fragment of CyaA under the control of a lac promoter | Karimova et al., 2005 |
| pUC19- FtsB-SD-FtsL | Ampicillin colE1 Ori | Encodes FtsB fused to T18 fragment of CyaA and FtsL under the control of a lac promoter | Karimova et al., 2005 |
| pUC19-FtsB | Ampicillin colE1 Ori | Encodes FtsB under the control of a lac promoter | This work |
| pUC19-FtsL | Ampicillin colE1 Ori | Encodes FtsL under the control of a lac promoter | This work |
| pUC19-FtsB-SD-FtsL | Ampicillin colE1 Ori | Encodes both FtsB and FtsL under the control of a lac promoter | This work |

**Table S6: Oligonucleotides for plasmid construction**

| Name | Forward primer (5' → 3') | Reverse primer (5' → 3') |
|---|---|---|
| Bd 1 | GATCCCATTAACGAACAGAGCGCGGATGTGCCGGGCGAAACCTTTTATCGCCTGGTGCCG-3' | GTACCGGCACCAGGCGATAAAAGGTTTCGCCCGGCACATCCGCGCTCTGTTCGTTAATGG |
| Bd2 | GATCCCATTAACGAACAGTGCGCGGATGTGCCGGGCGAATGCTTTTATCGCCTGGTGCC | GTACCGGCACCAGGCGATAAAAGGTTTCGCCCGGCACATCCGCGCTCTGTTCGTTAATGGTGTCT |
| Bd 3 | GATCCCATTAACGAACAGAGCGCGGATTGCCCAGGCTGCACCTTTTATCGCCTGGTTCCG | GTACCGGAACCAGGCGATAAAAGGTGCAGCCTGGGCAATCCGCGCTCTGTTCGTTAATGG |
| Bd4 | GATCCCACCGAAACCTTTTATCGCCTGGAAGATGGCGAATGGAGCCTGGTTCGCACCGTGGAAGCG | GTACCGCTTCCACGGTGCGAACCAGGCTCCATTCGCCATCTTCCAGGCGATAAAAGGTTTCGGTGG |
| Bd5 | GATCCGACCCGTTGGGAGCGTGTTCGCACCGAAGGCGACCGTACCTTTTATCGCCTGGTGGAACAG | GTACCTGTTCCACCAGGCGATAAAAGGTACGGTCGCCTTCGGTGCGAACACGCTCCCAACGGGTCG |
| Bd6 | GATCCGACCCGTTGGGAGCGTGTGCGCTGCGAAGGCGACCGCTGTTTCTATCGCCTGGTGGAGCCG | GTACCGGCTCCACCAGGCGATAGAAACAGCGGTCGCCTTCGCAGCGCACACGCTCCCAACGGGTCG |
| Bd7 | GATCCAAGCACCCGCAGTTGGGAAGCGGTGCGCCGCGAAGGTGATAAAACCTTTTATCGCCTGGTGGAAAACCAG | GTACCTGGTTTTCCACCAGGCGATAAAAGGTTTTATCACCTTCGCGGCGCACCGCTTCCCAACTGCGGGTGCTTG |
| Bd8 | GATCCCGCAGCGCGTTGGGAGGCGGTTCGCACCGAAGGCGACCGTACCTTTTATCGCCTGCTTCCACCGGAACCG | GTACCGGTTCCGGTGGAAGCAGGCGATAAAAGGTACGGTCGCCTTCGGTGCGAACCGCCTCCCAACGCGCTGCGG |
| Bd5 (pUTM18C) | GGATCCGACCCGTTGGGAGCGTGTTCGCACCGAAGGCGACCGTACCTTTTATCGCCTGGTGGAATAACCG | AATTCGGTTATTCCACCAGGCGATAAAAGGTACGGTCGCCTTCGGTGCGAACACGCTCCCAACGGGTCG |
| Bd8 (pUTM18C) | GATCCCGCAGCGCGTTGGGAGGCGGTTCGCACCGAAGGCGACCGTACCTTTTATCGCCTGCTTCCACCGGAATAACCG | AATTCGGTTATTCCGGTGGAAGCAGGCGATAAAAGGTACGGTCGCCTTCGGTGCGAACCGCCTCCCAACGCGCTGCGG |
| Optimized peptides (pUTM18C) | GAATCTGTATTTCCAGGGCGCTAGAGGGTCGACTCTAGAGGATCC | CGCTGTCATCATTTGTACATCTAGAATTCTTACTCATGACTTAGGTACC |
| Bd51-7 | GATCCCGCTAGCAAACGCTGGGTACTGGTGCGTACCCAGGGCAACGTGTCCTTTTATCGCTTAGTACCGTAAG | AATTCTTACGGTACTAAGCGATAAAAGGACACGTTGCCCTGGGTACGCACCAGTACCCAGCGTTTGCTAGCGG |
| $FtsQ_{49\text{-}276}$ | CCCTCTAGAGATGGAAGATGCGCAACGCC | GGGGAATTCTCATTGTTGTTCTGCC |
| $FtsQ_{49\text{-}246}$ | CCCTCTAGAGATGGAAGATGCGCAACGCC | CCCGAATTCTCAATCAACGTAGCTAATCCGTTTGCC |

| | | |
|---|---|---|
| FtsQ$_{P.A65\text{-}285}$ | CATGGATCCACGCGGCGCCGAGTACATCCTG | CTAGGTACCTTACTGCACGGCGCTGGCCGT |
| pETM11_rev_FtsQ$_{49}$ | GAATCTTTATTTTCAGGGCGCCATGGCGGCGATGGAAGATGCGCAACGCC | GAGCTCGAATTCGGATCCGGTACCACTTCATTGTTGTTCTGCCTGTGCC |
| T18-FtsQ | ACCTAAGTAAGTAAGAATTCCGATACCGAGCTCCAGTGAGCGCAACGCAATT | GCAACTGTTGGGAAGGGCGATTCATTGTTGTTCTGCCTGTGCCT |
| pUC19oligo | AGCTTGCGCTGCAGGCGG | GATCCCGCCTGCAGCGCA |

**Table S7: genes sequences**

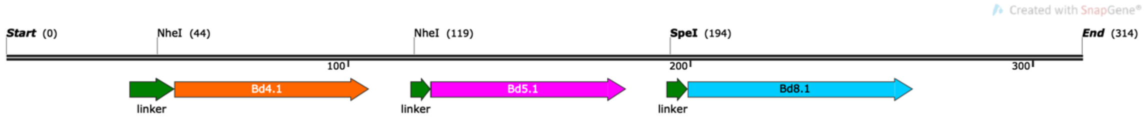


**Binder 458.1 (4.1-5.1-8.1)**
314 bp

DNA fragments have been ordered to clone the optimized peptides and the peptides targeting FtsQ of P.aeruginosa in pKT25 vector. To facilitate gene cloning, each DNA fragment was encoding for 3 peptides in a row with the first one in frame once cloned in the vector (fig above). To put the sequence encoding for the second peptide in frame, the built plasmid was digested by NheI while it was digested by both NheI and SpeI to put the third one in frame.

| **Name** | **Peptides encoded** | **DNA sequences** (5'→ 3') |
|---|---|---|
| Bd458.1 | Bd4.1<br>Bd5.1<br>Bd8.1 | GGCGCGCACGCGGCGGGCTGCAGGGTCGACTCTAGAGGATCCCGCTAGCAC<br>CGTGAGCTTTTATCGCCTGGTGGATGGCAAATGGGTGCTGGTGCGCACCGTGC<br>CGTAATAAGCATGCGCTAGCGAGCGCTGGGTACTGGTGCGTACCGAAGGCGA<br>TGTGTCCTTTTATGAATTAGTACCGTAATCCGGATAAACTAGTGCGGCGCAGTGG<br>GAACTGGTGAGCACCGAAGGCGATGTGAGCCTGTATCGCGAACTGCCGGCGC<br>CGTAAGGTACCTAAGTCATGAGTAAGAATTCACTGGCCGTCGTTTTACAAC |
| Bd458.2 | Bd4.2<br>Bd5.2<br>Bd8.2 | GGCGCGCACGCGGCGGGCTGCAGGGTCGACTCTAGAGGATCCCGCTAGCAC<br>CGTGAGCTTTTATCGCCTGGTGGATGGCAAATGGGTGCATGTGCGCACCGTGC<br>CGTAATAAGCATGCGCTAGCAGCCGCTGGGTGAAAGTGCGCACCGAAGGCGA<br>TGTGACCTTTTATGAACTGGTGCCGTAATCCGGATAAACTAGTAAACCGCAGTG<br>GGTGCTGGTGAGCACCGAAGGCGATGTGAGCTTTTATGTGGAACAGCCGGCG<br>CCGTAAGGTACCTAAGTCATGAGTAAGAATTCACTGGCCGTCGTTTTACAAC |
| Bd458.3 | Bd4.3<br>Bd5.3<br>Bd8.3 | GGCGCGCACGCGGCGGGCTGCAGGGTCGACTCTAGAGGATCCCGCTAGCAC<br>CGTGAGCTTTTATCGCCTGGAAAACGGCAAATGGCGCCTGGTGAAAACCATTC<br>CGTAATAAGCATGCGCTAGCACCCGCTGGGTGCATGTGCGCACCGAAGGCGA<br>TGTGACCTTTTATGAACAGGTGCCGTAATCCGGATAAACTAGTAAGGCCAAGTG |

| | | |
|---|---|---|
| | | GGTGCTGGTGAGAACCGAGGGCGACGTGAGCTACTACGAGGAGCAGCCCGC<br>CCCCTAAGGTACCTAAGTCATGAGTAAGAATTCACTGGCCGTCGTTTTACAAC |
| Bd458.4 | Bd4.4<br>Bd5.4<br>Bd8.4 | GGCGCGCACGCGGCGGGCTGCAGGGTCGACTCTAGAGGATCCCGCTAGCAC<br>CGTGAGCTTTTATGAACTGCGCGATGGCGAATGGCGCCTGGTGCGCACCGTG<br>CCGTAATAAGCATGCGCTAGCACCCGCTGGGTGCTGGTGCGCACCGAAGGC<br>GATGTGAGCTATTATGAAGAAGTGCCGTAATCCGGATAAACTAGTAAACCGCAG<br>TGGGTGCTGGTGAGCACCGAAGGCGATGTGAGCCATTATGTGGAACAGCCGG<br>CGCCGTAAGGTACCTAAGTCATGAGTAAGAATTCACTGGCCGTCGTTTTACAAC |
| Bd458.5 | Bd4.5<br>Bd5.5<br>Bd8.5 | GGCGCGCACGCGGCGGGCTGCAGGGTCGACTCTAGAGGATCCCGCTAGCAC<br>CGTGAGCTTTTATGAACTGGTGGATGGCGAATGGCGCCTGGTGCGCACCGTGC<br>CGTAATAAGCATGCGCTAGCACCCGCTGGGTGCTGGTGAGCACCGAAGGCGA<br>TGTGAGCTTTTATGAAGAAGTGCCGTAATCCGGATAAACTAGTAAACCGCAGTG<br>GCGCCTGGTGCGCACCGAAGGCGATGTGAGCTTTTATGAAGAACAGCCGGCG<br>CCGTAAGGTACCTAAGTCATGAGTAAGAATTCACTGGCCGTCGTTTTACAAC |
| Bd458.6 | Bd4.6<br>Bd5.6<br>Bd8.6 | GGCGCGCACGCGGCGGGCTGCAGGGTCGACTCTAGAGGATCCCGCTAGCAC<br>CGTGACCTTTTATCGCCTGGTGGATGGCGAATGGGTGCATGTGCGCACCGTGC<br>CGTAATAAGCATGCGCTAGCACCGAATGGCGCCTGGTGAGCACCGAAGGCGA<br>TGTGAGCTTTTATGAAGAAGTGCCGTAATCCGGATAAACTAGTGCGGCGCAGTG<br>GGAACTGGTGGAAACCCGCGGCGATGAAAGCCTGTATCGCGAACTGCCGGC<br>GCCGTAAGGTACCTAAGTCATGAGTAAGAATTCACTGGCCGTCGTTTTACAAC |
| Bd458.7 | Bd4.7<br>Bd5.7<br>Bd8.7 | GGCGCGCACGCGGCGGGCTGCAGGGTCGACTCTAGAGGATCCCGCTAGCAC<br>CGTGAGCTTTTATCGCCTGGTGGATGGCGAATGGGTGCATGTGCGCACCGTGC<br>CGTAATAAGCATGCGCTAGCAGCGTGTGGGTGAAAGTGCGCACCGAAGGCGA<br>TGTGACCTTTTATGAAGAAGTGCCGTAATCCGGATAAACTAGTAAAGCGCAGTG<br>GCGCCTGGTGCGCACCGAAGGCGATGTGAGCCTGTATGAAGAACTGCCGGC<br>GCCGTAAGGTACCTAAGTCATGAGTAAGAATTCACTGGCCGTCGTTTTACAAC |
| Bd458.8 | Bd4.8<br>Bd5.8<br>Bd8.8 | GGCGCGCACGCGGCGGGCTGCAGGGTCGACTCTAGAGGATCCCGCTAGCAC<br>CGTGAGCTTCTACAGACTGGAGGACGGCGAGTGGGTGCTGGTGAGCACCGTG<br>CCCTAATAAGCATGCGCTAGCAGCCGCTGGGTGCATGTGCGCACCGAAGGCG<br>ATGTGACCTTTTATCGCGAAGTGCCGTAATCCGGATAAACTAGTAAACCGCAGT |

| | | |
|---|---|---|
| | | GGGAACTGGTGCGCACCGAAGGCGATGTGAGCCTGTATCGCGAACTGCCGG<br>CGCCGTAAGGTACCTAAGTCATGAGTAAGAATTCACTGGCCGTCGTTTTACAAC |
| Bd458.9 | Bd4.9<br>Bd5.9<br>Bd8.9 | GGCGCGCACGCGGCGGGCTGCAGGGTCGACTCTAGAGGATCCCGCTAGCAC<br>CGTGAGCTTTTATGAACTGGTGGATGGCGAATGGCGCCTGGTGAGCACCGTGC<br>CGTAATAAGCATGCGCTAGCACCGTGTGGGTGAAAGTGAGCACCGAAGGCGAT<br>GTGACCTTTTATGAAGAAGTGCCGTAATCCGGATAAACTAGTAAAGCGCAGTGG<br>GAACTGGTGCGCACCGAAGGCGATGTGAGCCTGTATCGCGAACTGCCGGCG<br>CCGTAAGGTACCTAAGTCATGAGTAAGAATTCACTGGCCGTCGTTTTACAAC |
| Bd458.10 | Bd4.10<br>Bd5.10<br>Bd8.10 | GGCGCGCACGCGGCGGGCTGCAGGGTCGACTCTAGAGGATCCCGCTAGCAC<br>CGTGAGCTTTTATGAACTGCGCGATGGCGAATGGCGCCTGGTGAGCACCGTGC<br>CGTAATAAGCATGCGCTAGCACCCGCTGGGTGCTGGTGAGCACCGAAGGCGA<br>TGAAAGCTTTTATGAAGAAGTGCCGTAATCCGGATAAACTAGTAAACCGCAGTG<br>GCGCCTGGTGAGCACCGAAGGCGATGTGAGCCTGTATGAAGAACTGCCGGCG<br>CCGTAAGGTACCTAAGTCATGAGTAAGAATTCACTGGCCGTCGTTTTACAAC |
| BdP168$_{F15A}$ | BdP1<br>BdP6<br>Bd8$_{F15A}$ | GGCGCGCACGCGGCGGGCTGCAGGGTCGACTCTAGAGGATCCCGCTAGCC<br>GCGAAACCCTGTATCAGCTGGTGGATGGCGAAGTGGTGGCGAGCCCGCTGTA<br>ATAAGCATGCGCTAGCATTACCACCTATCTGAGCCTGGATGAACCGGAAGAAG<br>AACTGCTGGCGAAAGTGGCGGCGGCGCTGGCGGCGGTGGCGGCGGCGTAAT<br>CCGGATAAACTAGTGCGGCGCGCTGGGAAGCGGTGCGCACCGAAGGCGATC<br>GCACCGCGTATCGCCTGCTGCCGCCGGAATAAGGTACCTAAGTCATGAGTAAG<br>AATTCACTGGCCGTCGTTTTACAAC |
| BdP278$_{Y16A}$ | BdP2<br>BdP7<br>Bd8$_{Y16A}$ | GGCGCGCACGCGGCGGGCTGCAGGGTCGACTCTAGAGGATCCCGCTAGCAT<br>GGAAACCCTGTATCAGCTGCAACCGGATGGCACCGTGGTGGCGAGCCCGCT<br>GTAATAAGCATGCGCTAGCAAAACCACCTATTATGATGAACTGGGCCTGGAATA<br>AGTGAACCTAGCGTAATCCGGATAAACTAGTGCGGCGCGCTGGGAAGCGGTG<br>CGCACCGAAGGCGATCGCACCTTTGCGCGCCTGCTGCCGCCGGAATAAGGTA<br>CCTAAGTCATGAGTAAGAATTCACTGGCCGTCGTTTTACAAC |
| BdP388$_{AA}$ | BdP3<br>BdP8<br>Bd8$_{AA}$ | GGCGCGCACGCGGCGGGCTGCAGGGTCGACTCTAGAGGATCCCGCTAGCAA<br>AGAAACCCTGTATCAGCTGAACCCGGATGGCACCGTGACCGCGAGCGAATAA<br>TAAGCATGCGCTAGCATGGAATATATTACCGAAGAACGCGCGGAAGAACTGGA<br>ATAAGTGGTAGCGTAATCCGGATAAACTAGTGCGGCGCGCTGGGAAGCGGTG |

| | | |
|---|---|---|
| | | CGCACCGAAGGCGATCGCACCGCGGCGCGCCTGCTGCCGCCGGAATAAGG<br>TACCTAAGTCATGAGTAAGAATTCACTGGCCGTCGTTTTACAAC |
| BdP49 | BdP4<br>BdP9 | GGCGCGCACGCGGCGGGCTGCAGGGTCGACTCTAGAGGATCCCGCTAGCAA<br>AGAAACCCTGTATCAGTATAACGAAGAAACCGGCGAAGTGACCGCGAGCCCG<br>CTGTAATAAGCATGCGCTAGCAGCGTGATTGATGAAGCGGTGGAACGCATTAAA<br>AAAGAAATTGCGGAACGCGCGGCGAAATAATCCGGATAAACTAGTGCGGCGC<br>GCTGGGAAGCGGTGCGCACCGAAGGCGATCGCACCTTTTATCGCTGCCTGCC<br>GCCGGAATAAGGTACCTAAGTCATGAGTAAGAATTCACTGGCCGTCGTTTTACA<br>AC |
| BdP510 | BdP5<br>BdP10 | GGCGCGCACGCGGCGGGCTGCAGGGTCGACTCTAGAGGATCCCGCTAGCAT<br>GGAAACCCTGTATCAGCTGGAAGGCGATGAAGTGGTGGCGAGCCCGCTGTAAT<br>AAGCATGCGCTAGCGCGAAATATGTGGAAGCGCTGATTACCGAAGAAAACGCG<br>GAAGAAGCGGTGGCGGCGGCGAAAGCGGCGGTGAAATAATCCGGATAAACTA<br>GTGCGACCCGCTGGGAACGCGTGCGCACCGAAGGCGATCGCACCTTTTATCG<br>CTGCGTGGAATAAGGTACCTAAGTCATGAGTAAGAATTCACTGGCCGTCGTTTTA<br>CAAC |
